# High-Throughput Optimization of Paper-Based Cell-Free Biosensors

**DOI:** 10.1101/2024.10.03.616554

**Authors:** David C. Garcia, John P. Davies, Charles E. Davidson, Daniel A. Phillips, Aleksandr E. Miklos, Matthew W. Lux

**Affiliations:** US Army DEVCOM Chemical Biological Center, Aberdeen Proving Ground, MD, USA; Science and Technology Corporation, Belcamp, MD, USA

**Keywords:** Cell-free expression, synthetic biology, biosensors, paper-based sensors, shelf-stability, design of experiments

## Abstract

Cell-free expression systems maintain core cellular processes without intact cells and offer attractive properties as point-of-need biosensors. The ability to lyophilize, store, and use on-demand make these sensors usable in the field, and the lack of membranes means that there are no analyte transport issues and that new sensors can be deployed by simply adding a different DNA molecule. The lack of membranes also means that sensor designs and reaction optimizations can be screened in high throughput. While shelf stability has been demonstrated in specific cases using additives, these approaches are not universal to the myriad cell-free expression methods and formats. Here, we present new high-throughput screening methods to optimize cell-free expression formulations when embedded into paper for use as sensors. Our method leverages acoustic liquid handling to dispense reactions onto 384-well paper ticket formats and machine vision to quantify reaction performance from a colorimetric reporter enzyme. The throughput enabled shifts the bottleneck from experimental execution to selecting the experiments to execute; we therefore implement design-of-experiments to optimize the information gained from each design-build-test-learn cycle. We used these approaches to first optimize the performance of a low-cost cell-free expression formulation that was initially non-functional when embedded in paper, then further optimize it for tolerance to exposure to heat. With only 2 rounds of experimentation lasting 4 days total for each goal, the result are an energy mixture with 8% of the materials cost of a commonly used version and a formulation of excipients that maintain 60% of activity after 6 hours of storage at 50 °C and. Finally, we showcase the use of the cost-optimized formulation in a 3D-printed paperfluidic device where it outperforms the standard formulation at much lower cost.

---

Synthetic biology’s increasing use of large datasets has significantly improved the process of repurposing and reengineering natural biological systems towards addressing global challenges. For instance, rewiring molecular biosensors and their associated genetic networks have resulted in simple-to-use and cheap biological diagnostics capable of responding to environmental perturbations dangerous to human health. The use of cell-free expression (CFE) systems enables gene expression from non-living, *in vitro* biochemical reactions powered by lysed cells and/or purified transcription and translation components combined with mixtures of function-encoding nucleic acids, cofactors, and energy sources^1,2^. Biological sensors developed from these CFE systems have led to a multipurpose platform capable of sensing a variety of analytes including: pathogen biomarkers, environmental contaminants, and industrial precursors and products ^3–8^.

Extending the capability of cell-free biosensors requires that they can effectively be produced, stored, and used in austere and resource-limited environments^11^. The ability to freeze-dry and embed cell-free biological components into matrices promises to enable their field-deployability^9,10^. Cell-free paper sensors address fieldability needs as they can be easy to read without external devices when paired with visual reporters, are small and disposable, and can be adapted to different applications by changing the DNA encoding the sensor functionality^12^. However, cell-free sensors can be rendered ineffective due to non-ideal conditions such as breaks in cold-chain and reduced access to relevant components^13^. Previous efforts have demonstrated methods involving cryoprotectants that enable shelf stability of CFE reactions for months at room temperature and 37°C in non-paper formats ^14–16^; one early study showed ~20% activity after 1 year of storage of CFE reactions on paper without cryoprotectants^4^. The cost of CFE reactions is also an important factor for the technology to reach its potential. While several studies have assessed materials costs, one formulation offers substantially lower cost while maintaining substantial yield^17^; until recently, to our knowledge no further published efforts had taken advantage of this formulation, and still none have tested it for biological sensors. Beyond cost, disruptions to supply lines due to the COVID19 pandemic impacted the cell-free community as critical components, especially tRNAs, became increasingly scarce to the point of near absence from traditional marketplaces^18,19^. As a result, developing a low-cost CFE formulation that works well on paper and is more resilient to supply chain disruptions would have a substantial impact on the viability of CFE sensors for a range of applications.

Addressing this need benefits from the use of high-throughput testing platforms and methodologies capable of significantly reducing the timeframe for design-build-test-learn (DBTL) cycles^20^. Previous work using high-throughput methods for aqueous reactions have been shown to effectively improve CFE performance for new organisms, metabolic reactions, and general CFE reaction dynamics^21–23^. None of these methods have been applied to sensors or reactions involving solid matrices like paper; instead, paper sensors have been tested using time- and labor-intensive methodologies, limiting their use as a testing platform. To alleviate this bottleneck, we developed a high-throughput testing platform specifically for reactions on paper and implemented a design-of-experiments (DOE) approach to take advantage of the throughput. DOE allows for systematic exploration and optimization of complex systems by elucidating factors with a significant effect on a desired response; this allows experimenters to determine optimum formulations in a complex system using a fraction of the number of experiments compared to testing one factor at a time or even some machine learning methodologies^20,23–25^. We applied these methods to optimize a low-cost CFE system that was originally inactive on paper to perform on par with standard systems, then optimized additives to improve the stability of CFE sensors to storage at elevated temperatures. With only 2 rounds of experimentation, one exploratory and one predictive, our efforts led to cell-free paper ticket sensors that maintained performance at 8% of the original cost as well as formulations capable of functioning following exposure to 50 °C temperatures for 6 hours. Further, the decreased-cost mixture was found to be more effective than standard mixtures when used in a 3D-printed paperfluidic device.

## Materials and Methods

### 384-well Ticket Preparation

Wax tickets were printed directly on 1CHR chromatography paper (Whatman) using a Xerox ColorQube printer and prepared for use in the following manner: each ticket was baked at 125 °C for 5 minutes, allowed to cool at room temperature for 5 min, incubated for 1 h in 5% BSA, washed thrice with diH20, and allowed to dry overnight in a fume hood. Paper tickets were stored in fume hood until use.

### Preparing and Running Cell-Free Reactions

All CFE reactions utilized either a pY71 plasmid with a T7-expressed LacZ insert (pY71-LacZ) or a combination of two plasmids using the pUCGA vector, one for T7-expression of trH RNA and the other for T7-expression of swH-lacZ RNA. All DNA was ordered from Aldevron as Giga Preps.

### T7 Polymerase Expression Protocol

An overnight culture of cells was grown in LB Media with Ampicillin at 37 °C. The next day 500 mL of LB was inoculated with 1 mL of the overnight culture and grown at 37 °C until the OD600 reached 0.4. Protein expression was induced by adding IPTG to a final concentration of 100 μM, and the culture was incubated overnight at 37°C. The cells were then pelleted at 5,000 g for 20 minutes at 4 °C and resuspended in 25 mL lysis buffer (1/2x PBS, 0.01% Triton X-100, 1 mM EDTA). The cells were lysed by sonication, and the lysate was centrifuged to pellet the cellular debris at 30,000 g for 30 minutes at 4 °C. The supernatant was then transferred to a clean 50 mL tube, with 1.0 mL of Ni-NTA purification resin and incubated for 1 hour at 4 °C with shaking. We used 20 mL of wash buffer to clean the resin (50 mM phosphate, 300 mM NaCl, 10 mM imidazole, pH 7.5) and eluted using a gravity flow (50 mM phosphate, 300 mM NaCl, 250 mM imidazole, pH 7.5). Protein-containing fractions were combined and dialyzed in a 3.5k MWCO dialysis cassette against 2L of S30 buffer overnight at 4°C and subsequently at −20 °C.

### E. coli Lysate Preparation

Lysate preparation was carried out using E. coli Rosetta (DE3) ΔlacZ strains for all experiments. A 100L culture of Rosetta (DE3) ΔlacZ cells was processed for lysate production similar to the previously described production of BL21(DE3)* lysate, but with modifications to accommodate production at scale. Briefly, 750 mL starter cultures (1.5 L total) were grown for 16 hours at 37°C with 200 rpm shaking incubation. Prior to inoculation, 100 L of 2X YT+P culture media supplemented with 5 mL of antifoam 204 (Sigma, A8311) in an IF 150L (New Brunswick Scientific) fermenter was allowed to aerate overnight with a rotor speed of 100 rpm and 20 standard liters per minute (slpm) airflow at 37°C. Following inoculation to a starting OD600 of 0.05, the fermenter settings were adjusted to 300 rpm, 50 splm, and the dissolved oxygen (DO) was calibrated to 100%. Upon reaching an OD600 of 0.6-1.0, the culture was induced with a final concentration of 1 mM isopropyl B-D-1-thiogalatopyranoside (IPTG) (GoldBio, I2481C). Once DO reached 50%, the rotor speed was increased to 500 rpm. At an OD600 of 3.5, the culture was cooled to 4°C, centrifuged in a prechilled Powerfuge pilot, 1.1 L bowl system (CARR Biosystems) within approximately 8 hours, and the pelleted bacteria was subsequently processed as described previously.

### Preparing and Running Cell-Free Reactions

All CFE reactions utilized either a pY71 plasmid with a T7-expressed LacZ insert (pY71-LacZ) or a combination of two plasmids using the pUCGA vector, one for T7-expression of trH RNA and the other for T7-expression of swH-lacZ RNA. All DNA was ordered from Aldevron as Giga Preps. CFE reactions contained 30% v/v lysate and PANOx-sp or Cai buffer as described in detail previously^25^. All components and final concentrations are summarized in Table S1–3. After mixing, CFPS reactions were lyophilized in plates, tubes, or vials depending on the scale described for specific experiments. Base CFE reactions were modified with excipients and reagents using working solutions based on each experiment and distributed by Echo liquid handler; PANOx-SP, Cai energy mixture, and excipient working solutions found in **(Tables S4–5)**. Amounts of additional reagents for the optimized formulations are described in the main text, and Supplementary Data. Paper tickets were prepared for liquid dispensing by directly attaching using adhesive (3M Super 77 Multipurpose Spray Adhesive) to a 384-well Thermofisher microwell plate (Thermofisher Catalog Number: 142761). Reaction components were dispensed directly onto the 384-well ticket using an Echo 525 acoustic liquid handler (Beckman Coulter) or multipipettor for volumes below 1000 nL and above 1000 nL, respectively. Unless otherwise noted all swH/trH tickets were prepared to contain a final concentration of 50 ng/µL pUCGA-trH-lacZ per reaction when resuspended in 2uL final volume. The Echo Plate Reformat software and custom scripts were used to prepare Echo transfer protocols. Reactions were lyophilized using a lyophilizer (VirTis AdVantage XL-70, SP Scientific) using a bell jar attachment. The shelf was prechilled to −40 °C, and the condenser to between −65 and −70 °C. The paper tickets were frozen for at least 5 min in a −80 °C freezer. After removing the plate, it was immediately placed in the lyophilizer, the vacuum was activated, and the shelf temperature set to −20 °C overnight. The 384-well tickets were prepared for reading by attaching the ticket via adhesive directly to a 384-well plate that had been filled with 30 µL of water per well. Ticket spots were resuspended in 2 µL of 50 ng/µL pUCGA-trH-lacZ DNA in diH20 using a multipipettor unless otherwise noted. Following rehydration, the plates were sealed with a plate cover from a microwell plate (Thermofisher Catalog Number: 142761), the edges wrapped in parafilm to prevent evaporation, and placed directly into a scanner for imaging (see Data Collection and Machine Vision Analysis section).

### DOE and Statistical Analysis

All statistical analyses in this section were performed using Stat-Ease Design-Expert 13 and SAS JMP® Pro 15 software. To optimize the Cai reaction mixture, I-Optimal criterion was used to create optimal Mixture-Amount DOEs based on a quadratic statistical starting model. The design used a starting model that was quadratic on the mixture side and quadratic on the process side, with a “Kowalski-Cornell-Vining” (KCV) structure that restricted the crossing of the mixture and process term (amount) to 2-way. The design is capable of fitting non-linear blending effects up to second order. Within each of the 14 or 16 individual components for adding Cai or PANOx-SP components, respectively, the low and high limits varied by component. Additional details of the DOE models are available in the **Supplementary Results** and **Supplementary Data File**.

For the heat tolerance optimization mixtures, I-Optimal criterion was used to create an optimal Mixture DOE based on the quadratic statistical starting model. The Mixture type DOE design was selected in order to maximize experimental efficiency^30^. A quadratic starting model having the ability to characterize all linear blending effects for each of the individual 14 formulation components as well as all possible non-linear blending effects involving any pair of the 14 formulation components was selected. For heat tolerance reactions, a formulation the 14 individual components were limited to a range of 0 to 1000 nL. Additionally, each sample formulation was subject to a total volume of 3000 nL.

Functional Data Analysis (FDA) for both case studies was applied via the “Functional Data Explorer” platform within SAS JMP® Pro 15 software. A Functional Principle Components (FPC) decomposition was applied to the response curves, decomposing each curve into a Functional Principle Component “1” (FPC1) and Functional Principle Component “2” (FPC2) score where the FPC1 and FPC2 scores for an individual sample response curve represent the deviation from the overall mean response curve. A statistical model for the DOE generated was then fit to the FPC1 and FPC2 responses. Forward and backward step regression using the Akaike Information Criterion (AiC) was used to reduce the model for the FPC1 and FPC2 response, respectively. The R-Square Predicted metric was used along with the “check-points blends” (in-line validation samples) to validate the DOE based prediction models’ predictive capability. Optimization was performed using the Stat-Ease Design-Expert 13 Numerical Optimization feature in parallel with the SAS JMP® Pro 15 software Prediction Profiler Platform. Both softwares start with the DOE based prediction model and, after converting the FPC response to a zero to one desirability scale, use numerical methods to find the formulations that best optimize the response.

### Data Collection and Machine Vision Analysis

Image processing from 384-well tickets was performed using three major steps. The ticket is placed on a flatbed scanner (Epson, Perfection V600 Photo) housed within an incubator to produce a time course of sensor images at 37 °C. Each plate is arranged on the scanner at an arbitrary point and scanned directly following rehydration. Custom software was used to capture raw 16-bit color images of the scanner bed at specified intervals. The series of images were then analyzed to monitor the biosensor’s response over time using scripts that process the images as follows: time course images are placed in consistent orientation; spots are delineated from the image background and segmented into individual positions **(Figure S1)**; average RGB value for each spot tile is computed; and the color change (ΔE) and hue values (ΔH) returned for each well within the context of CIELAB perceptual color space. For the purposes of this work, an arbitrary point of visibility was designated at ΔE=10 wherein experimenters agreed the color shift was very clear (**Figure S2**).

### 3D-Printed Sensor Design

All designed parts were created in SolidWorks (SolidWorks) computer aided design software and exported as .stl files for slicing within the Ultimaker Cura software (Ultimaker). “Engineering – Normal” parameters were selected on Cura utilizing support for overhangs greater than 45° and without additionally printed build plate adhesion. Parts were printed using Ultimaker ABS and Ultimaker Breakaway Support filament on the Ultimaker S5 dual extrusion 3D printer. Prior to printing, the glass bed was pre-treated with AirWolf ABS adhesion solution prior to preheating the build plate to 125 °C. All paper layers were printed and prepared as described above prior to the addition of the cell-free extract mix. The agarose-topped layer was prepared by directly adding 5 µL of 1.25% molten agarose to each paper spot. Each extract layer was prepared as described above. Both the cell-free extract layer and the agarose-topped layer were then frozen at −80 °C and lyophilized as described above. Sensors were assembled by placing each layer on the sampling plate, adding a form fitted “quad” top plate over the agarose-topped layer, and covering the top with MicroAmp Clear Adhesive Film (Thermo Fisher). Sensors were tested through the addition of 25 µL of 50 ng/µL pUCGA-trH into the sample chamber and incubating in the flatbed scanner at 37 °C.

## Results and Discussion

### High-Throughput Testing Platform of Paper-embedded Reactions

Due to the labor-intensive nature of paper-based cell-free biosensor development, we reasoned that the development of a high-throughput testing platform to explore large combinatorial search spaces would accelerate improvement of their overall function. Paper-based reactions in the literature are typically prepared using biopsy punches to cut out small discs that are placed into microtiter plates or wax-printed paper tickets with modest numbers of wells; reactions are then spotted, lyophilized, and rehydrated, typically by hand (**Figure 1A**)^26,27^. While this method effectively creates portable sensors, it is not amenable to high-throughput experiments as even simple titrations of fixed reagents create a high-experimental load. For example, a more complex experiment using the excipients in this study to improve durability of a sensor to heat creates a combinatorial search space of 6,103,515,625 potential combinations when using only 5 fixed concentrations (**Figure 1B**). To improve cell-free paper sensors testing throughput, we replicated the commonly used 384-well format using wax-printing on paper, yielding a 127.76×85.47 mm 384-well paper ticket template with 3.5 mm diameter wells. Biologically inactive adhesive was used to apply the paper ticket over a standard 384-well plate, and an Echo 525 Acoustic Liquid Handler was used to transfer reaction components from an Echo source-plate to the sensor wells using standard Echo protocols. The tickets were then lyophilized, rehydrated using DNA, and CFE reaction activity measured by color change using a flatbed scanner and custom machine vision and image analysis software, described in detail in the methods, that extracts a measure of human-perceived color change (ΔE) (**Figure 1C**, **Figure S1**).

**Figure 1:**
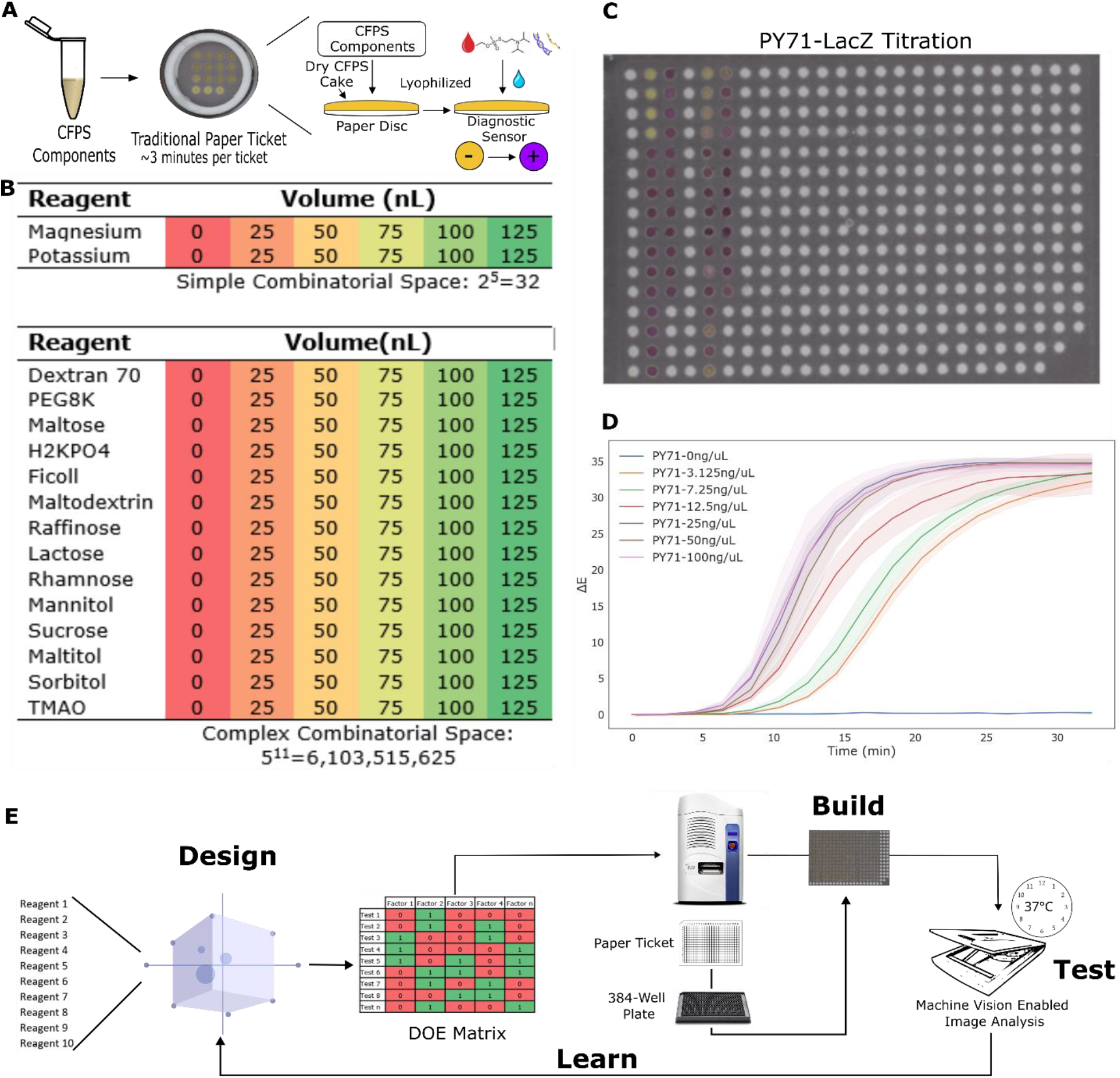
Optimizing paper-based cell-free reactions in high throughput. **A.** Cell-free components are lyophilized directly to a paper disc or ticket and rehydrated with an aqueous analyte to start protein production, in this case LacZ (β-galactosidase), to affect a color change from yellow to purple by cleavage of the CPRG substrate. **B.** An example experiment changing fixed concentrations of salts affecting CFE **(top)** and a more complex experiment assessing protective additives with 5 fixed concentrations **(bottom)**. **C.** Representative image of 384-well wax-printed paper ticket. **D.** Traces of ΔE from titrations of a plasmid encoding for constitutively expressed LacZ (pY71-LacZ) in the PANOx-SP CFE system extracted from time course images of a ticket. **E.** Graphic illustration of the DBTL cycle used to explore the combinatorial space of cell-free paper-ticket compositions. Initial tests of excipients and reagents defined an initial DOE matrix that was tested and the resulting data used to define a predictive model for active and optimal reagent concentrations.

Figure 1C shows a representative 384-well ticket with reactions expressing LacZ (β-galactosidase), an eye-readable colorimetric reporter that has been used in a wide range of applications^28^. As an initial validation of the approach, we performed a titration curve of a plasmid (pY71-LacZ) that constitutively expresses LacZ from the strong T7 promoter. For these initial tests, the PANOx-SP formulation was used. The reactions were dispersed onto a 384-well ticket as a master mix with only the plasmid dispersed separately; the ticket was then lyophilized and rehydrated using water. The titration resulted in increasing rates of LacZ production commensurate with the concentration of the DNA transferred, plateauing at 25 ng/µL, which is consistent with prior work indicating a saturation of translational machinery in the CFE reaction (Figure 1D)^29^.

To explore the vast combinatorial space inherent to non-fixed quanta we decided to employ exploratory DOE with an objective function geared towards improving the rate of our cell-free reactions (Figure 1D). The first day of our DBTL cycle is initiated with an exploratory DOE using a list of excipients and additives using volumes ranging from 25 to 1000 nL. The reaction compositions in the exploratory DOE are transferred to the Echo, dispensed on to a 384-well paper ticket, and lyophilized overnight. On the second day, the samples are rehydrated, placed onto a flatbed scanner, color change data extracted, and the resulting data fit to the DOE model.

The loop is then restarted with optimized reaction conditions predicted by the model to be run with replicates. DOE offers the ability to efficiently and simultaneously characterize the influence that each formulation component has on the response of interest while at the same time testing for interactions between formulation components and process factors. This is a distinct advantage of DOE experimentation over “one factor at a time” experimentation as we are able to reduce the multidimensionality of the problem to a manageable number of potential test cases. The method allows for large amounts of data to be gathered, analyzed, and fitted to an empirical statistical model that can serve as a prediction for a response at any possible combination of formulation components within the entire design.

### Optimization of a Low-Cost CFE Mix

As a model diagnostic sensor for the remainder of this work, we used a lyophilized CFE reaction containing a toehold switch plasmid (pUCGA-swH-lacZ) that expresses LacZ in the presence of a target trigger RNA. The biosensor reactions are rehydrated by the addition of a plasmid that constitutively expresses the cognate trigger RNA (pUCGA-trH) (Figure 2A). Lyophilized tickets with identical spots containing 50 ng/µL of pUCGA-swH-lacZ were activated using varying levels of pUCGA-trH (**Figure S3**). We further tested our ability to dispense reagents by measuring the transfer and subsequent effect of more viscous reagents. RNASE inhibitor has been noted as being important to the function of cell-free paper-based biosensors due to the natural presence of nucleases on the matrix^10,30^. We titrated the concentration of RNASE inhibitor both to measure the ideal concentration of a critical component and to assess the platform’s ability to modulate a viscous reagent. RNASE inhibitor Murine dissolved in 50% glycerol was titrated onto identical CFE reactions, lyophilized, and resuspended in 50 ng/µL of both pUCGA-swH-lacZ and pUCGA-trH. Increasing concentrations of RNASE inhibitor improved the rate of the reaction, with doubling the typical concentration of RNASE inhibitor to 2.4 units/µL providing the best reaction rate and a loss of benefit at 4.8 units/µL (**Figure S4**).

**Figure 2:**
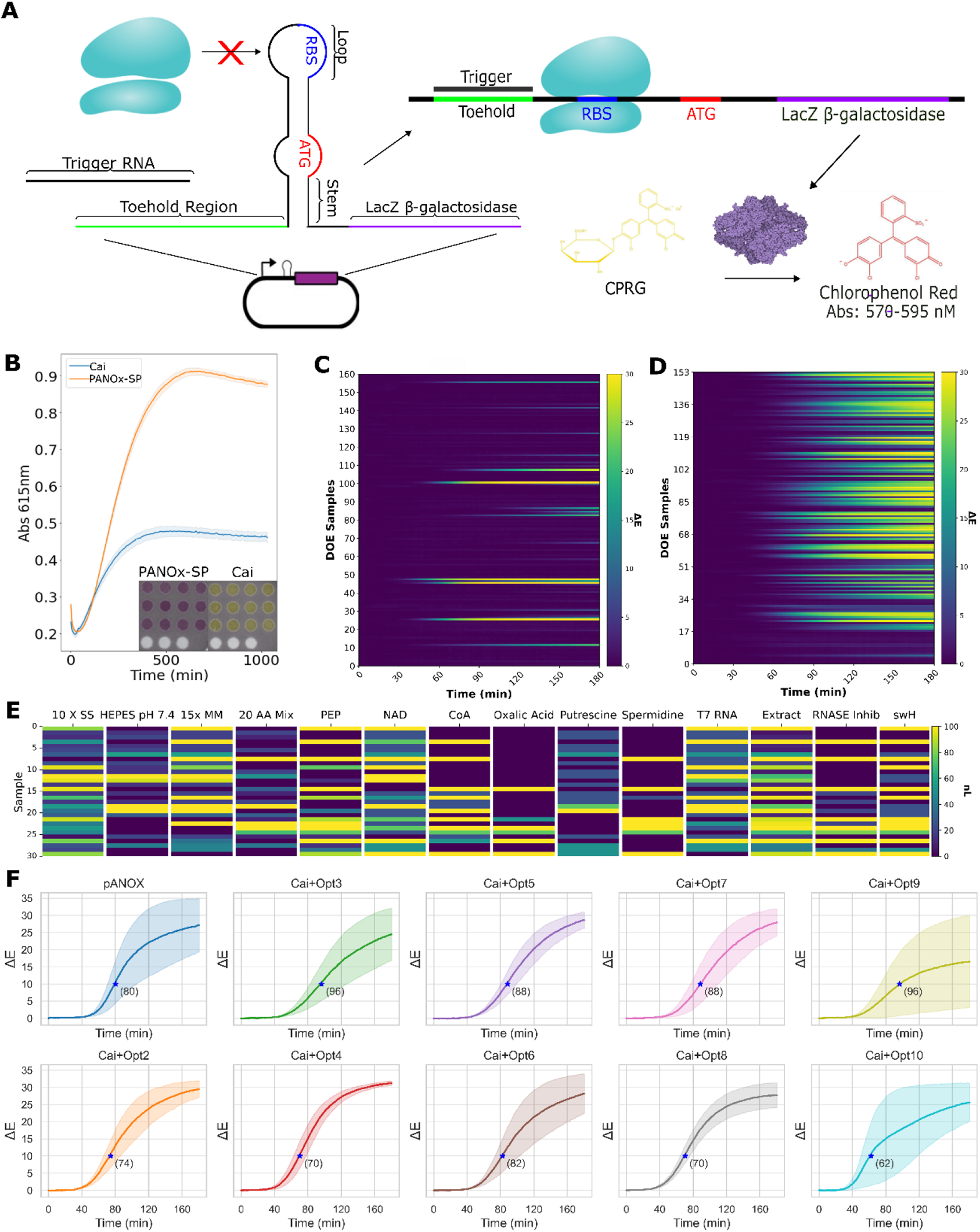
Optimization of low-cost CFE reactions for performance in paper-based biosensor reactions. **A.** The presence of trigger RNA disrupts the stem-loop structure of the switch RNA and allows for translation of LacZ (β-galactosidase). **B**. Comparison of PANOx-SP and Cai CFE biosensor reactions in a liquid format in plates; **(inset)** the same reaction components in the paper-based biosensor format after a 3 h incubation **C.** DOE heatmap produced from the addition of Cai reaction components to the baseline Cai CFE formulation. **D.** Similar experiment was performed using PANOx-SP reaction components added to the baseline Cai formulation. **E.** Heatmap of compositions of the 30 optimal mixtures predicted from DOE analysis. **F.** Time courses of paper tickets using predicted optimized mixtures adding PANOx-SP reagents to baseline Cai mixture. Blue stars indicate the point at which the paper ticket reaction reached ΔE≥10. Standard deviations shown as shadows on each trace were derived from triplicate reactions.

A minimized version of the Cytomim CFE system as reported on by Cai and colleagues demonstrated the ability to drop the materials cost of a CFE system by an order of magnitude through the removal of non-essential and expensive components^17^. At a total cost of ~$445/L, the reaction mixture published by Cai heretofore referred to as the Cai reaction mixture is significantly cheaper than the ~$5,570/L cost of the traditionally used PANOx-SP mixture (**Table S1**)^29^. This is relevant both for the purposes of making the biosensors affordable in the context of low-resource point-of-need applications and to limit the need for components such as tRNAs, nucleotide triphosphates, and volatile energy sources such as PEP that can be difficult to acquire when supply chains are disrupted. However, we found Cai CFE reactions to produce no clear color change when used with our paper-based biosensors, despite functioning in liquid reactions (Figure 2B). We sought to resolve this limitation as well as reduce the cost per test by optimizing a cheap but inactive-on-paper cell-free energy mixture to function with paper sensors.

We hypothesized that the absence or low concentration of one or more critical components resultant from the cost-minimization efforts was a likely cause of the inability of the Cai to function on paper. To test this hypothesis, we designed a DOE framework to add combinations of components of the Cai formulation by dispensing them directly on 384-well tickets containing normal Cai reactions (Methods). The tickets were then lyophilized to both to normalize the final reaction volumes and because lyophilization is an important aspect of real-world applications of the technology. After rehydration, color change traces were extracted as previously. Though our machine vision approach is capable of quantifying small shifts in color, a threshold of ΔE≥10 was used to note a point at which the reactions became obviously visible by eye and served as a barometer for sensor function for the remainder of this study (**Figure S2**). We prepared an initial 16-component DOE using the components of the standard Cai mix, adding variable amounts of each component in volumes ranging from 0-1000 nL (Figure 2C; **Table S5**). While some combinations of added reagents did produce detectable signal, the data was not sufficient to fit the DOE model and make predictions of optimal formulations. Following these results, we speculated that some component(s) of the PANOx-SP mix might explain the poor relative performance of the Cai mix compared to PANOx-SP on paper tickets. We next used a 153-sample framework adding variable amounts of 14 PANOx-SP components to baseline Cai reactions (**Table S4**). Many of these combinations showed a marked improvement in the function of the Cai mix on paper sensors (Figure 2D). Additional details about the DOE, data processing, and results are available in the **Supplementary Results**, **Figures S5-7**, and Tables S7-8.

Our initial DOE model generated 30 optimal reaction mixtures that spanned diverse combinations of added components (Figure 2E). We test the top 10 predicted optimals and each reached the threshold for visibility within 1.5 hrs, except for Cai-Opt1 which was inactive (data not shown) (Figure 2F). Nine formulations were active and with statistically no difference in speed when compared to the PANOx-SP controls (**Table S9**). These formulations further had costs ranging from $0.0013-$1.96 per 1 µL reaction added to $0.00044 for baseline Cai; these costs range from ~43-fold cheaper to 35-fold more expensive than the $0.056 for PANOx-SP, with higher costs driven almost entirely by additional DNA (**Table S10**). Here we predicted optimal formulations purely based on performance; it is likely that these costs could be driven down further by instead predicting formulations that maximize performance per unit cost.

Analyzing the combinations of additives that result in high performing formulations do not provide simple, obvious intuition into the determinants of improved performance. Notably, all 14 components appeared at both the highest and lowest levels in at least one of the 30 predicted optimal formulations, though some do appear more frequently. By considering the blending coefficients for the first and second order effects predicted by the model to have the greatest impact, it is clear that none of the components in isolation explain the improvements in performance (**Table S7**). The second order effects offer scant additional insight and instead highlight the complexity of the system (**Table S8**). For example, spermidine appears in 4 of the 10 strongest estimated second order effects both as a negative effector (with T7 RNAP and HEPES) and as a positive effector (with RNASE inhibitor and 20 AA Mix). It is worth pointing out that the DOE approach makes no attempt to model the complete system; instead, it aggregates estimates of first and second order effects in a way that can find improved performance after testing only a tiny fraction of a large search space. Moreover, interpretation of the effects of individual components is complicated by the Mixture-Amount design wherein constraints on the total amounts added mean that changing one variable necessitates changing others simultaneously. One noteworthy observation is that the model estimates that RNASE inhibitor provides a positive effect in combination with Spermidine, CoA, and PEP, but a small negative effect alone, suggesting that our earlier conclusion that additional RNASE inhibitor improves the reaction for the PANOx-SP system is dependent on one or more specific components in the mix. Ultimately, despite providing limited insights into the inner workings of the system, the model reliably predicted diverse, high-performing formulations. Until mechanistic models are developed that can fully capture the highly complex dynamics of CFE reactions, DOE offers a powerful tool to rapidly optimize formulations for a particular use case.

### Engineering Cell-Free Biosensor Heat Tolerance

The need for biosensors in austere environments inherently requires that the biological systems are capable of functioning outside ideal laboratory conditions. For instance, high salt, temperature, or pH environments can deactivate biological sensors. These problems can be magnified if the tools in question require notoriously difficult to maintain cold chains. We sought to improve the robustness of our sensors to elevated temperatures during storage through the addition of excipients with previously reported impacts on the function of lyophilized products following exposures to environmental stresses (**Table S6**).

During the sublimation process, the disruption of hydrogen bonding pairs can result in protein aggregation. To resolve this issue, excipients are often added to purified protein components in order to stabilize electrostatic complexes during drying. Additionally, exposure to temperatures above a protein’s melting temperature can unfold regions with thermodynamically unfavorable refolding and lead to aggregation and ultimately inactivity of the biosensor. Preliminary experiments with paper biosensors using the PANOx-SP system exposed to 6 hr incubation at 50 °C were found to be completely inactive (data not shown). We hypothesized that an effective combination of excipients to improve heat tolerance could be found by applying a DOE analysis to the lyophilization and heat testing process. We produced a DOE matrix testing combinations of 14 excipients shown to provide protective effects in other studies^15,32–35^ by adding them to the baseline PANOx-SP system and measuring the activity following heat challenges. An initial 6 hr incubation at 50 °C showed several reactions with a substantial improvement in the heat tolerance (Figure 3A).

**Figure 3.**
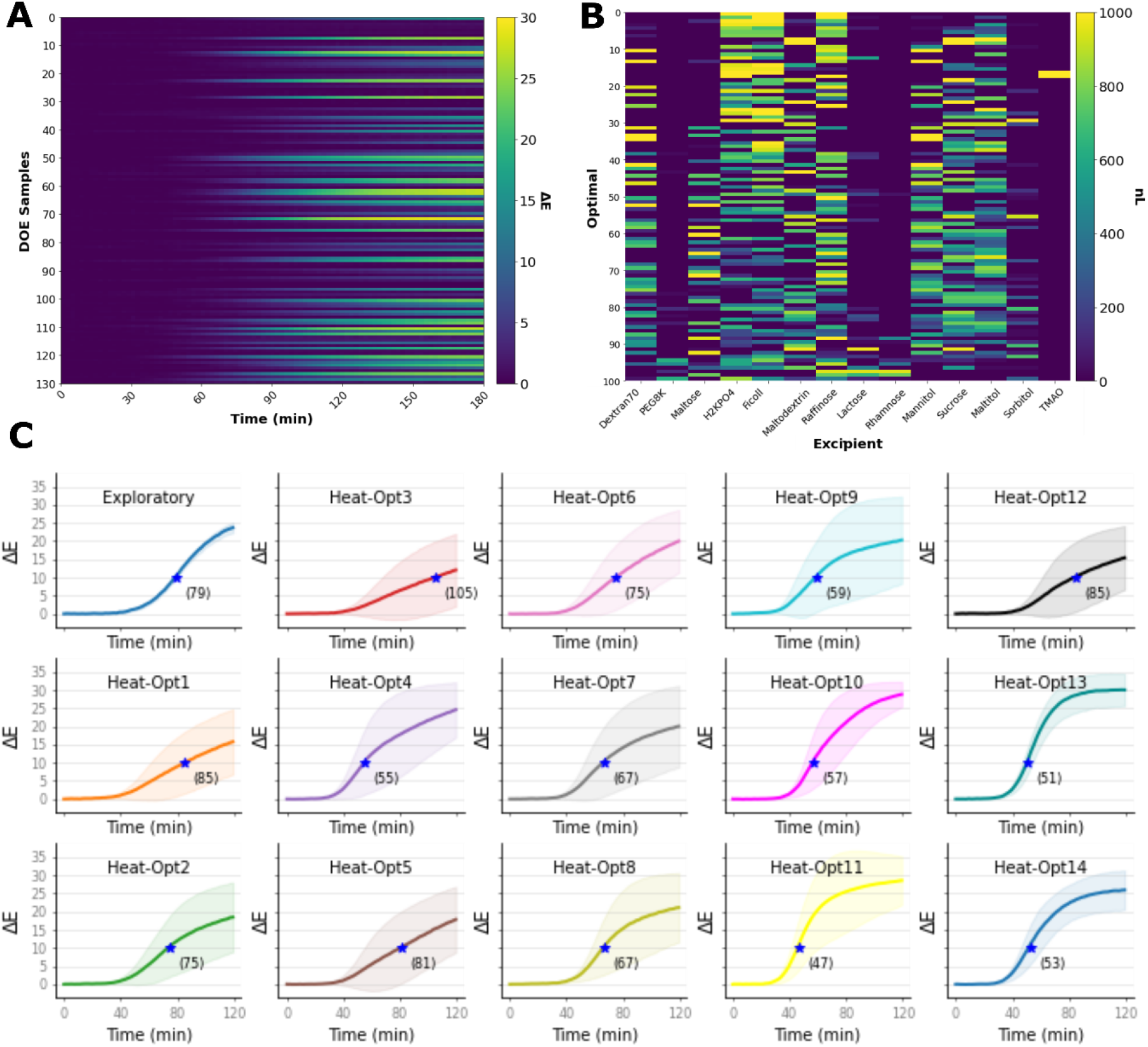
Cell-free biosensors were tested for improved activity following heat exposure using various excipients to improve performance. **A.** Heatmap of activity of DOE-generated combinations of lyophilization excipients to the baseline PANOx-SP CFE mix after exposure of lyophilized tickets to 50 °C for 6 hr. **B.** Heatmap of additive volumes for the 30 optimal mixtures predicted from DOE. **C.** Traces of predicted optimal mixtures following heat exposure. The exploratory graph shows the average time course of the active reactions from the exploratory experiment. The top 14 predicted optimal formulations were tested. Blue star indicates time to ΔE≥10. Standard deviations shown as shadows on each trace were derived from triplicate reactions.

Unlike the prior optimization, there were clear trends in the components that appeared in predicted optimal formulations: some excipients (e.g. PEG, lactose, and rhamnose) appeared rarely or not at all in the predicted top formulations, while others (e.g. H2KPO4, Ficoll, and Raffinose) appeared frequently (Figure 3B). Considering the linear and non-linear blending coefficients for the model, however, again yields a complicated picture (**Tables S11-12**). For example, while PEG, lactose, and rhamnose had relatively strong negative linear effects, PEG+rhamnose and lactose+rhamnose had relatively strong positive effects. This observation is explained by how the coefficients are weighted to predict the FPC1 value according to our Mixture-Amount constraints. More specifically, component amounts are pseudo-coded to range from 0 to 0.333 such that no single component can go above 1000 nL out of a maximum allowed 3000 nL total amount. For the example of PEG, lactose, rhamnose added at maximum amounts, the predicted FPC1 value is calculated as:

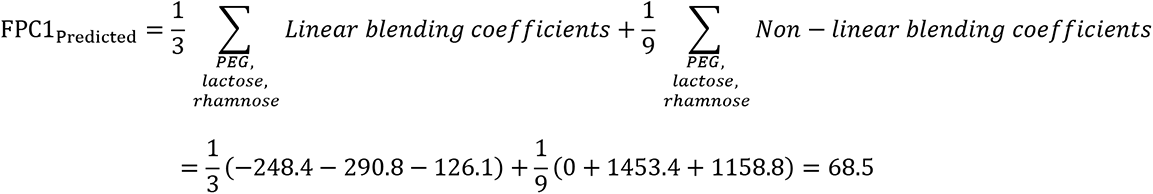

Since 68.5 is much low compared to the predicted optimals (e.g. #30 is predicted at 205.3), this apparent discrepancy is clarified. This observation can be interpreted as PEG, lactose, and rhamnose each having negative effects alone, with rhamnose having synergistic effects with both PEG and lactose that do not overcome the negative individual effects. While deeper insights into how these complex formulations perform would be highly welcome, as discussed above, the purpose of the DOE approach is to predict optimal formulations despite limited understanding of the system.

Fourteen predicted optimal formulations along with the top performing formulation from the exploratory DOE were further tested in quadruplicate by exposing the tickets to a 6 h incubation at 50 °C (Figure 3C). Notably, several of these reactions appeared to perform favorably to untreated PANOx-SP samples from the Cai optimization experiments (Figure 2F), though we did not include that control to enable a direct comparison. We nonetheless checked the statistical significance of these apparent improvements (run on different tickets on different days) and found no significant difference in time to reach a ΔE≥10 for any of the optimal formulations compared to PANOx-SP, which is unsurprising given substantial variability between replicates (**Table S11**; see discussion on variability below). While the Heat-Opt11 formulation reached the ΔE threshold for visibility at 47 min, the best reported in this study, it is not clear if the additives truly improved the performance of the base PANOx-SP. While we did not further explore the performance impact of these additives without heat exposure, other work has noted the dual purpose of maltodextrin as a lyoprotectant and energy source, suggesting that such improvements are possible.

Following our success optimizing the low-cost extract and improving the heat tolerance of our paper ticket sensors, we sought to merge the two-systems by evaluating the effect of adding the heat tolerance excipients optimized for the expensive PANOx-SP reaction mixture to our optimized Cai formulations. We chose 4 optimized Cai formulations (Cai-Opt3 and 9 that use minimal added DNA and Cai-Opt1 and 4 that add maximum DNA) and 2 optimized heat-tolerance formulations (Heat-Opt 11 and 13 for the fastest mean time to ΔE≥10) and exposed pairwise combinations of these formulations on tickets to 50 °C for 0, 3, or 6 h (Figure 4 and **Tables S9, S13**). We found that even without heat exposure, all combinations except the two involving Cai-Opt9 performed poorly but were still active; after heat exposure those combinations performed progressively worse, with four of the six combinations never reaching ΔE≥10 after 6 h exposure to 50 °C. Both combinations involving Cai-Opt9, however, performed similarly to the earlier PANOx-SP controls (mean time to ΔE≥10 of 80, 90, and 96 min for PANOx-SP, Heat11+Cai9, and Heat13+Cai9, respectively), and retaining some activity after heat exposure (mean time to ΔE≥10 ranging from 152 to 184 min). Further work would likely uncover combinations that provide better heat tolerance while maintaining low costs.

**Figure 4.**
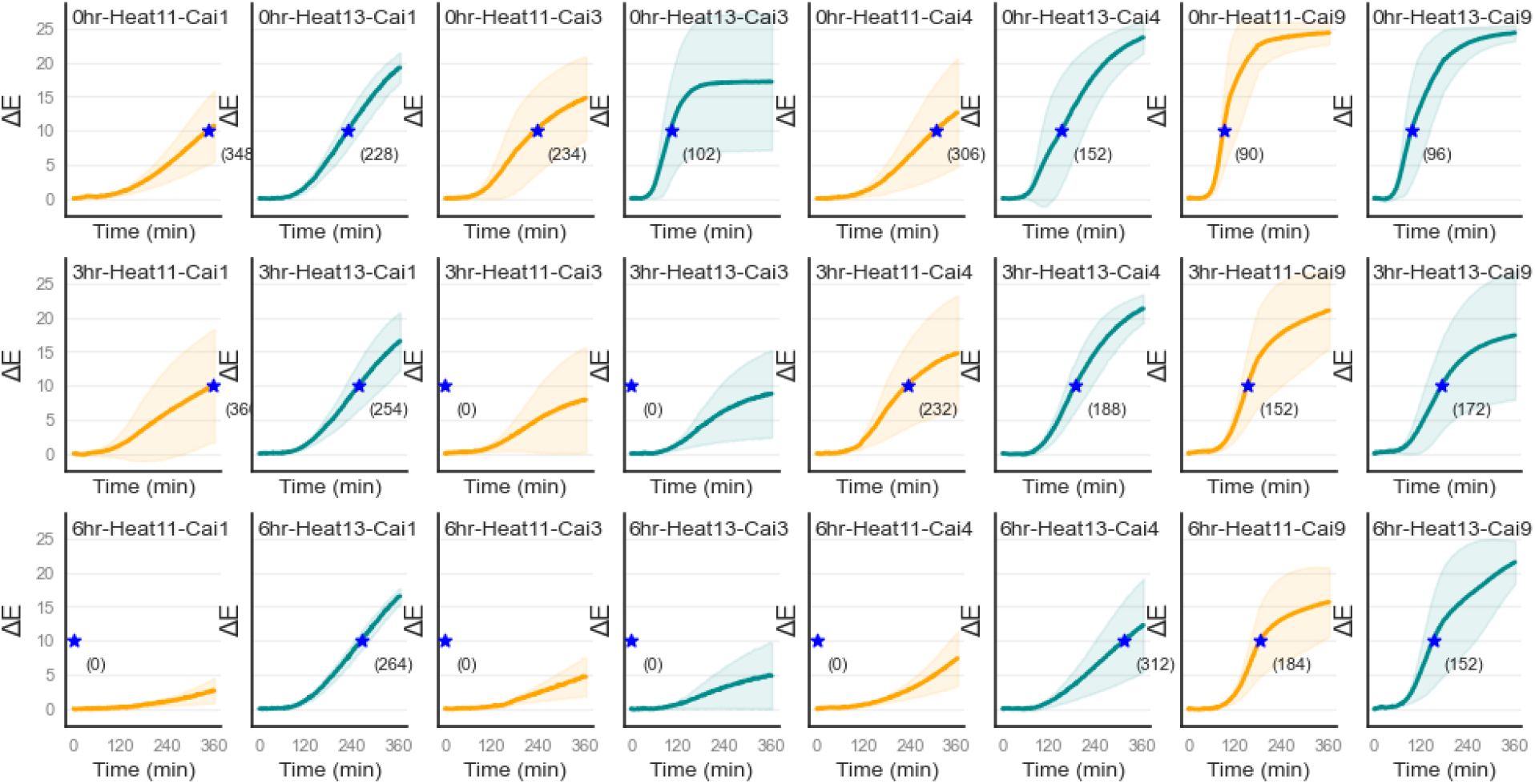
Heat tolerance of combinations of optimized Cai formulations and heat-tolerance excipient combinations optimized using PANOx-SP as the baseline. Combinations of Cai-Opt1, 3, 4, and 9 and Heat-Opt11 and 13 were added to baseline Cai on tickets and exposed to 50°C for 0, 3, or 6 h. Shaded error bars are one standard deviation from n=4 replicates. A blue star indicates the time at which the cell-free biosensor reached a ΔE of 10; if no replicates reached the threshold, a value of 0 is shown.

## Handheld Biosensors Effectively Make use of Optimized CFPS Mixtures

To showcase the effectiveness of our optimized reactions, we designed and created a handheld paperfluidic device to show the potential of the colorimetric sensor as a diagnostic tool. A 4-layer wax-printed paperfluidic sensor housed in a 3D-printed cassette (Figure 5A, **Figure S8**) was built using either the unoptimized Cai mix, PANOx-SP, or an optimized version of the Cai mix (Cai+Opt4) and tested by flowing pUCGA-trH plasmid through the device. As expected, the Cai mixture without optimization did not produce any detectable result, whereas both the PANOx-SP and the optimized Cai reaction mixtures were all visible within 42 minutes (Figure 5B-C). Interestingly, the optimized Cai mixtures led to a faster time to visibility than the standard PANOx-SP system (ΔE≥10 at 26 vs 42 mins, respectively; p=0.034), becoming statistically differentiable at 32 min (p<0.05). These reaction times outpace the 384-well ticket apparatus (ΔE≥10 at 80 vs 88 min for PANOx-SP and Cai-Opt4, respectively; Figure 2F), indicating that the handheld biosensor system improves the function of the CFE reaction both in general and differentially for the optimized formulation. Notable differences are sealing of the device to limit evaporation and the presence of an agarose-hydrogel layer intended to enhance flow in the paperfluidic system. Given that hydration of the reaction mixture can be crucial to function, both of these factors likely help to maintain an environment more amenable to the CFE reactions^31^. This observation shows the importance of the physical and environmental factors that can modify the function of cell-free biosensors and opens new avenues for further research and development. Rehydration, dilution of the components, and variable environmental humidity and temperature have all been shown to be critical factors, especially in the context of biological sensing^36^. This work further incentivizes context-specific modifications for biological sensing platforms as directly porting one reaction mixture to a different context can have unintended consequences.

**Figure 5.**
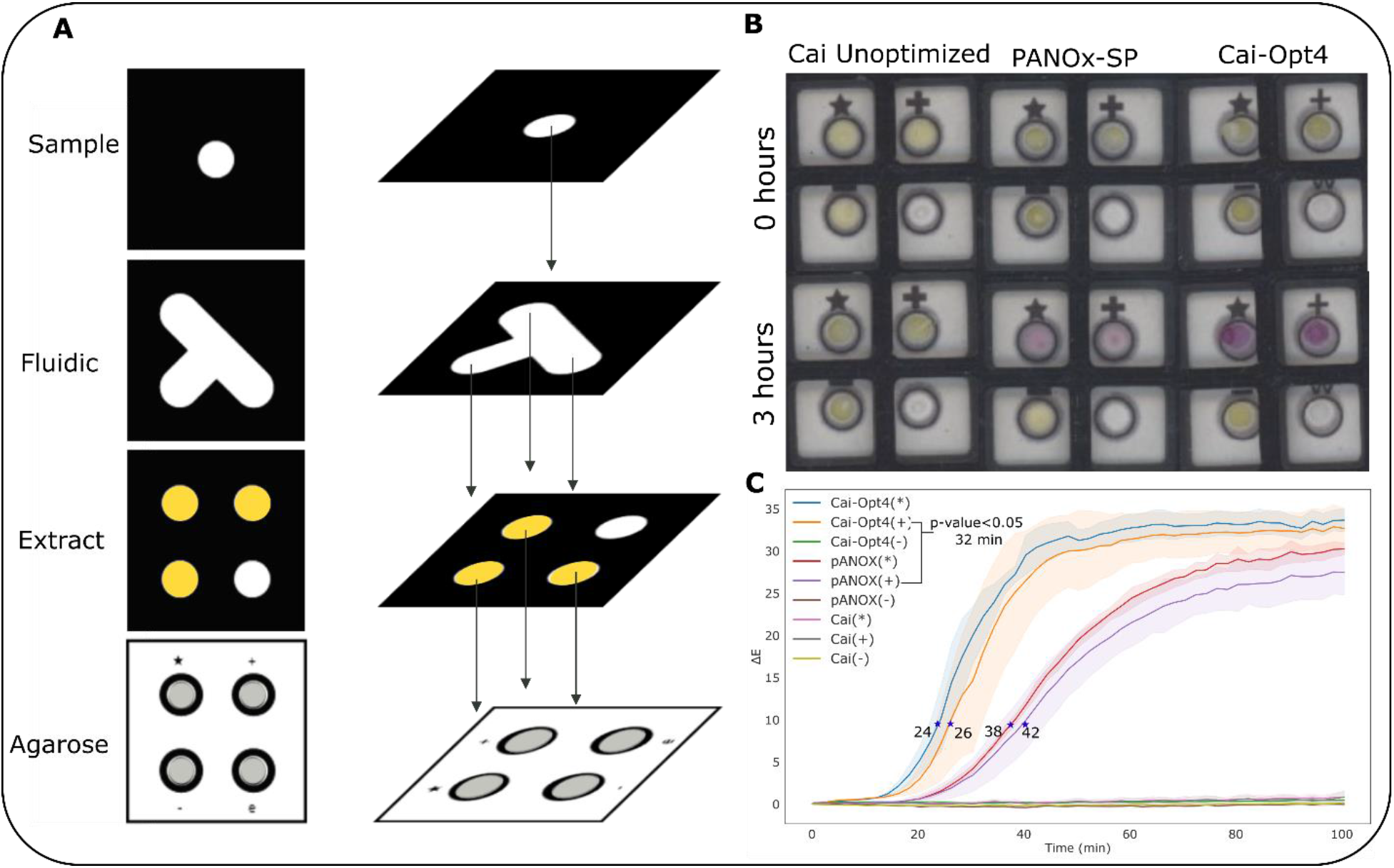
Performance of optimized biosensors in a fieldable paperfluidic device. **A.** Schematic of a wax-printed paperfluidic device wherein sample is added to the sample layer, disseminated using a paper fluidic layer to the individual reaction chambers containing cell-free extract, and visualized on an agarose-topped layer held together by a 3D printed holder (**Figure S8**). The “★” indicates a sample chamber with a CFE reaction and pUCGA-swH-lacZ; “+” indicates a positive control containing a CFE reaction with both pUCGA-swH-lacZ and pUCGA-trH plasmids; “-” indicates a negative control containing a CFE reaction with no DNA; and “e” being an empty node intended to identify any leaking between layers. Both the top and bottom layer 3D-printed enclosures are sealed with transparent film following sample addition. **B.** Assembled sensor devices were imaged and analyzed using the same scanning rig and software as above. **C.** Time courses of three replicate devices. The shadow indicates one standard deviation. Blue stars indicate time to cross ΔE≥10. One tailed T-test shows statistically significant difference compared to negative controls at 32 min for Cai-Opt4 and PANOx-SP sample reactions.

### Assessment of Variability

In the initial tests of our experimental platform titrating pY71-LacZ plasmid, we observed reasonably consistent error bars (Figure 1D). In later cases, however, we found that error bars for replicate experiments were larger, resulting in a lack of statistical significance between conditions even when differences in means appear large, e.g. the previously discussed differences between the heat-tolerance optimal formulations and PANOx-SP. We therefore assessed the CV of time to ΔE≥10 across all data sets (Figure 6). We found that the initial pY71-LacZ titrations were anomalously consistent (CV=6.9%) compared to other 384-well ticket experiments (CV=28.0-39.5%). Previous efforts to measure variability of CFE reactions has found that ~6-10% is typical for a single experimenter across days when prepared by hand, suggesting that the substantial increase in variability is a result of small volume dispensing errors either from the acoustic liquid handler or baseline master mixes. More careful calibration and validation of proper dispensing would be needed to validate this hypothesis and subsequently mitigate any issues. Other possibilities are that time to ΔE≥10 using an enzymatic reporter is more variable than the endpoint GFP fluorescence used in the other studies or that something about the paper ticket format increases variability; however, because in one case we saw similar consistency, we suspect dispensing errors are more likely to be the cause. While such variability limits fine tuning of reaction compositions, our results show that rapid, efficient screening through very large combinatorial design spaces is still viable. We note also that the average variability in our device format was 11.2%, indicating that optimal formulations can perform reasonably consistently in a more applied setting.

**Figure 6:**
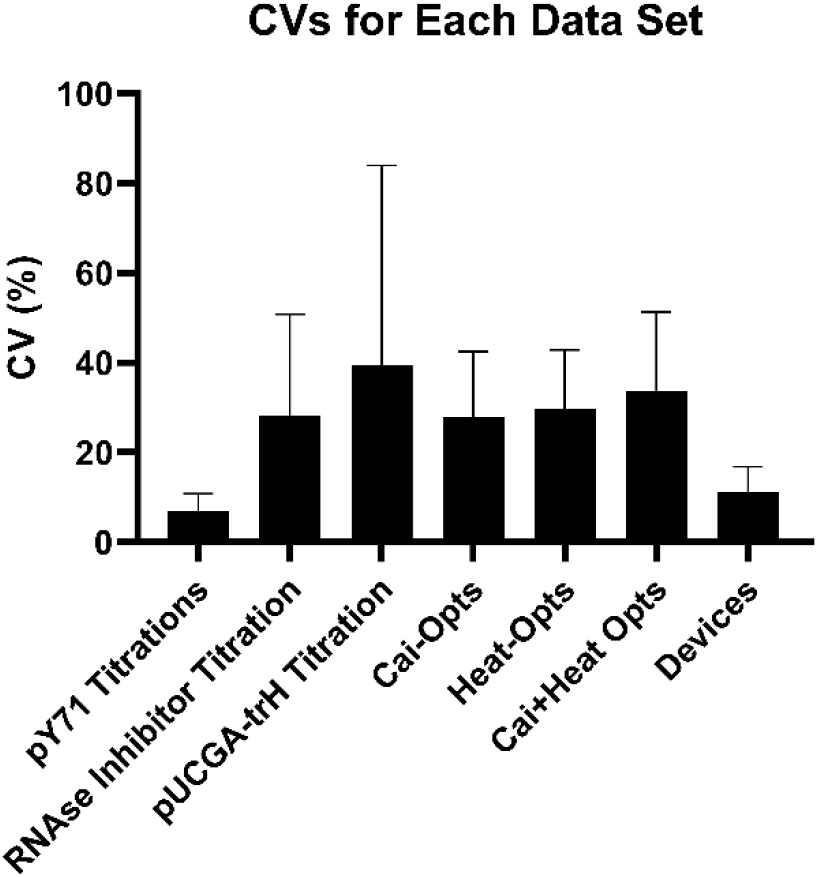
Variability within replicates across experiments in this study. Bars represent the average within-replicate CV across each experiment list. Error bars represent one standard deviation.

## Conclusions

Scaling the production of cell-free biological sensors requires cost-effective and stable methods of producing them as diagnostic tools. This work shows how high throughput experimentation driven by DOE can rapidly identify substantial improvements in performance at low costs and with tolerance to heat exposure in paper formats relevant to application. Given the complexity of these formulations, traditional optimizations varying one or two variables at a time are intractable; with the approaches outlined here, a single researcher can reduce enormous search spaces to a manageable number of experiments that can be executed in two consecutive days. The ability to then immediately port these improvements into a fieldable diagnostic device from low-cost, easy-to-produce components underscores the potential for such sensors to have real world impact. We anticipate that continued development of new CFE sensors paired with further improvements to the performance, cost, stability, and deployability of CFE reactions will lead to the technology achieving its potential impact.

## Acknowledgments

Funding was provided by a congressional program titled Cell-free Biomanufacturing. This project was supported in part by an appointment to the NRC Research Associateship Program at DEVCOM Chemical Biological Center for DCG administered by the Fellowships Office of the National Academies of Sciences, Engineering, and Medicine.

## Author Contributions

DCG: Conceptualization, Methodology, Investigation, Data curation, Writing – Original Draft; JPD: Methodology, Formal analysis; CED: Software; DAP: Methodology; AEM: Methodology, Supervision; MWL: Conceptualization, Writing - Review & Editing, Supervision, Project administration, Funding acquisition.

## Supplementary Results

### Design of experiments

#### Compressing data with Functional Principal Component analysis

In order to fit the DOE models described above, the curves were first decomposed into Functional Principal Components (FPC). Results of the FPC decomposition of DOE for adding PANOx-SP components to baseline Cai are described in this section. After an initial FPC decomposition, we plotted the FPC1 scalars to identify any systematic issues **(Figure S7)**. The experimental design is such that output should be randomly distributed across the samples, so any observed patterns indicate potential issues. Of the first 17 samples, 16 had minimal activity. We expect that this anomaly arose from a dispensing error by the acoustic liquid handler.

Because the anomaly was systematic, we excluded samples 1-17 from further analysis. This anomaly further raises the question of other potential dispensing errors that do not result in an obvious systematic error. While we do not rule out this possibility, we note that DOE methods are generally robust to experimental errors.

After removing the outliers, we found that 94.8% of the variation could be explained by a single Eigenfunction (FPC1) representing a generic sigmoid shape **(Figure S8)**. A second Eigenfunction (FPC2) explains an additional 4.85% of variation by dampening the slope of the response and introducing a positive slope to the plateau. Because FPC1 alone explains the data well, we used the FPC1 scalar as our fit metric.

An alternative choice to using FPC1 as the design objective would be to use the time until ΔE≥10. Since our goal is to create fast sensors, this alternative choice would be more direct; however, the use of FPC1 does indirectly select for response time by maximizing the linear scaling of FPC1 without discarding additional information about curve shape. To check what impacts the choice of a different objective function might have had, we re-ran the DOE analysis with time to ΔE≥10 as the objective. We found that the profile of median amount of each component added was similar across the top 10 predicted optimals, suggesting that the search space represented by each set of predicted optimals is likely similar **(Figure S9)**. Since little difference was observed, we chose to proceed with FPC1 as our objective function because of our anticipated-but-unverified notion that capturing the full curve information can help mitigate the impact of outliers.

#### Descriptions of DOE models used

The initial design for this experiment was a screening design intended to identify whether components of the Cai formulation could be added to recover activity of baseline Cai reactions. A quadratic Mixture Amount design with 16 components and a single process factor, labeled “Total Volume”, was used **(Supplementary Data Files)**. The process factor allowed for variation in the total number of droplets dispensed across all components added to the base Cai reaction. Because the reactions are then lyophilized, we do not expect this process factor to play a significant role in the predictive model. A Quadratic starting model was used with a “KVC” cross. This design is I-Optimal and capable of fitting non-linear blending effects up to second order and interactions with the “Total Volume” factor up to 2-way. This design models the influences of the relative ratios of the 16 components as well as the “Total Volume”. Despite identifying some combinations that yielded a response, the high number of non-responsive formulations resulted in an ability of the DOE model to fit the data using a backward (BiC) reduced statistical model.

The second attempt mirrored the first except 14 components from the PANOx-SP formulations were added instead, as described in the main text. The detailed design is available in the **Supplementary Data Files**. The resulting data was adequate for fitting the DOE model. A backward (BiC) reduced statistical model was fit to the FPC1 data. The resulting model (R^2^=0.6848, p<0.0001) had 34 terms including linear effects for the 14 reaction components and 20 additional non-linear blending effects that were influential enough to be included in the model. These effects and their blending coefficients are detailed in **Table S7–8**. These effects are interpreted in the main text.

The DOE for heat tolerance followed the same approach except 14 excipients were added to baseline PANOx-SP reactions. Details of the DOE is available in the **Supplementary Data** These effects and their blending coefficients are detailed in **Table S11-12**. These effects are again interpreted in the main text.

---

**Figure S1:**
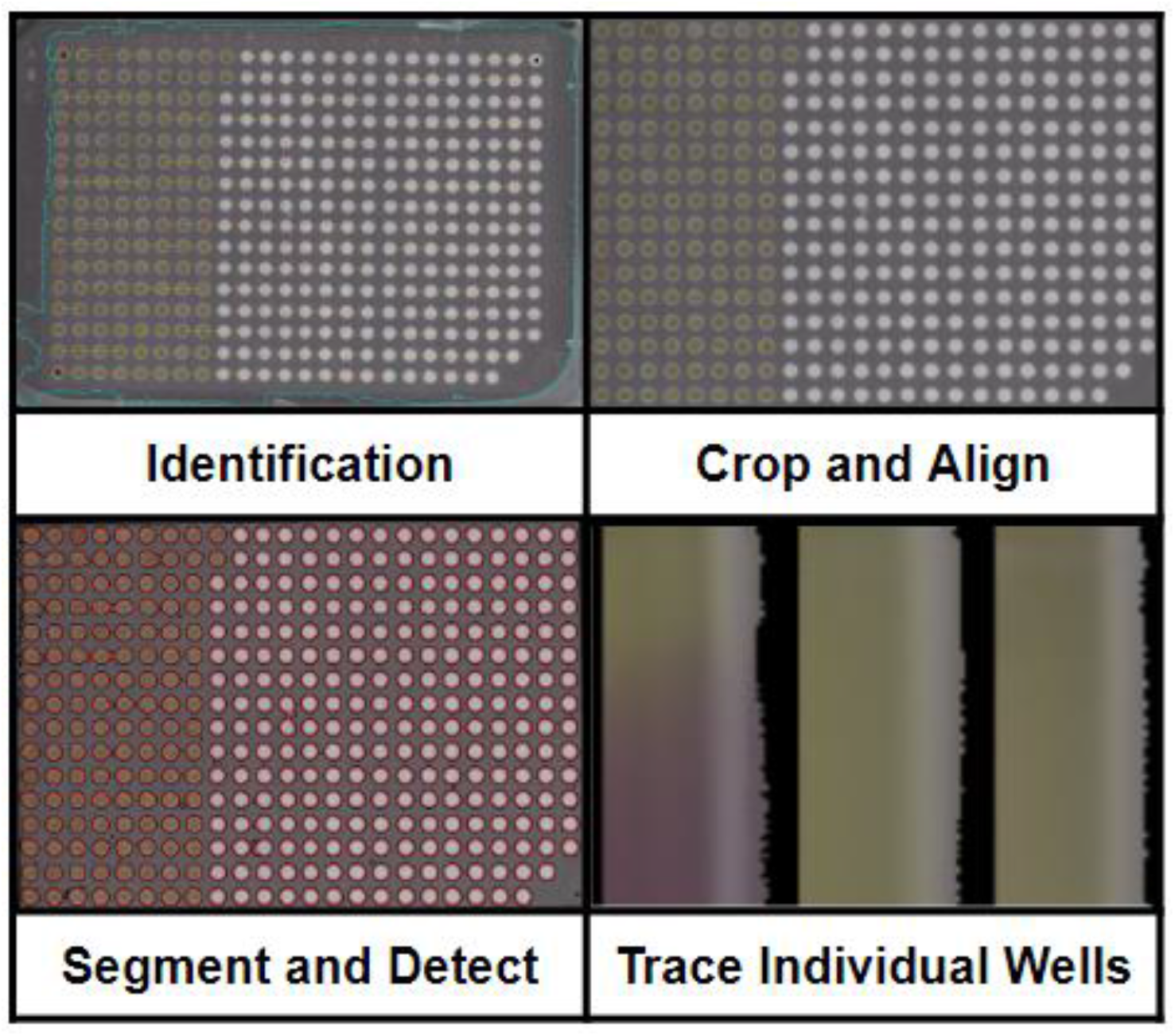
Representative example of 384-well CFE paper ticket analysis software extracting color change information for individual wells.

**Figure S2:**
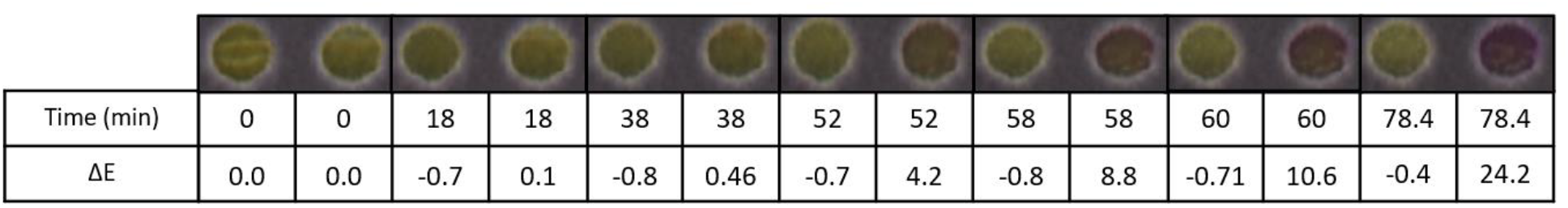
A representative image of the color change over time in a cell-free biosensor using a colorimetric reporter. Though a perceivable color change is seen earlier, we chose ΔE≥10 as a threshold at which the color change is obvious.

**Figure S3:**
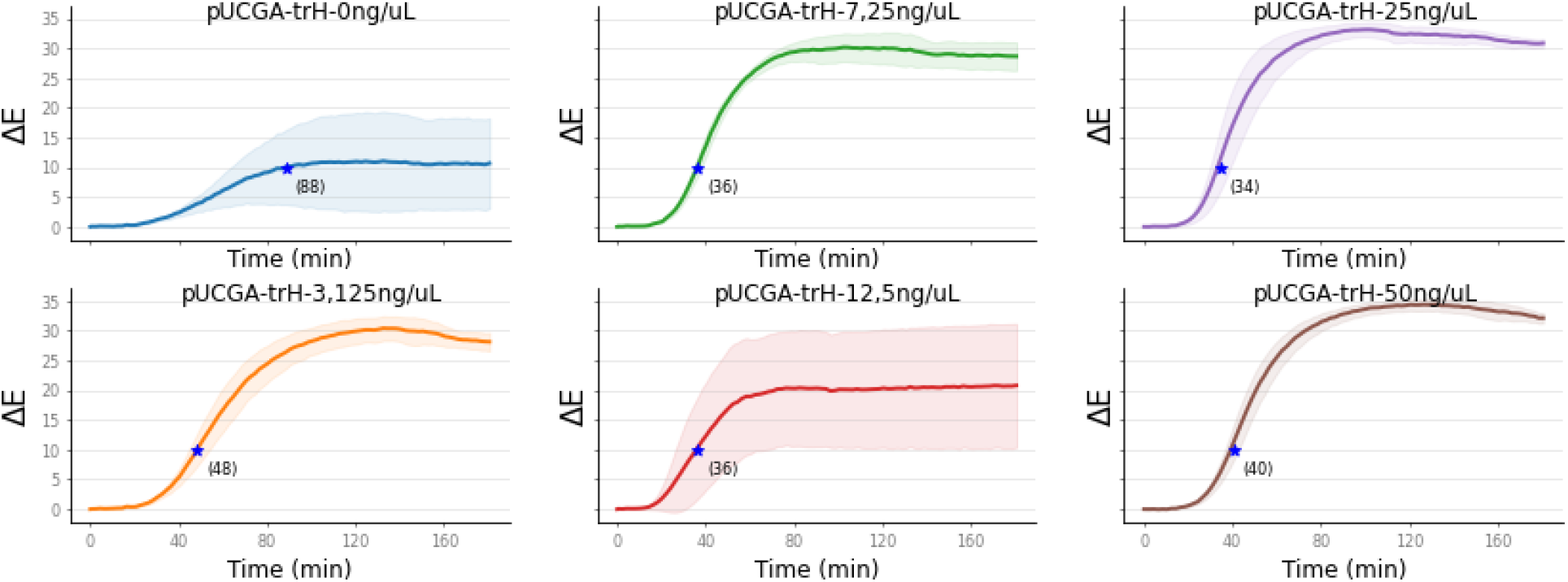
A PANOx-SP CFPS reagent mix was prepared with the same amount of pUCGA-swH-LacZ plasmid directly added before lyophilization and activated by resuspending using various concentrations of the commensurate trH. The blue star indicates the point at which a ΔE≥10 was reached. Standard deviations shown as shadows on each trace were derived from quadruplicate reactions.

**Figure S4:**
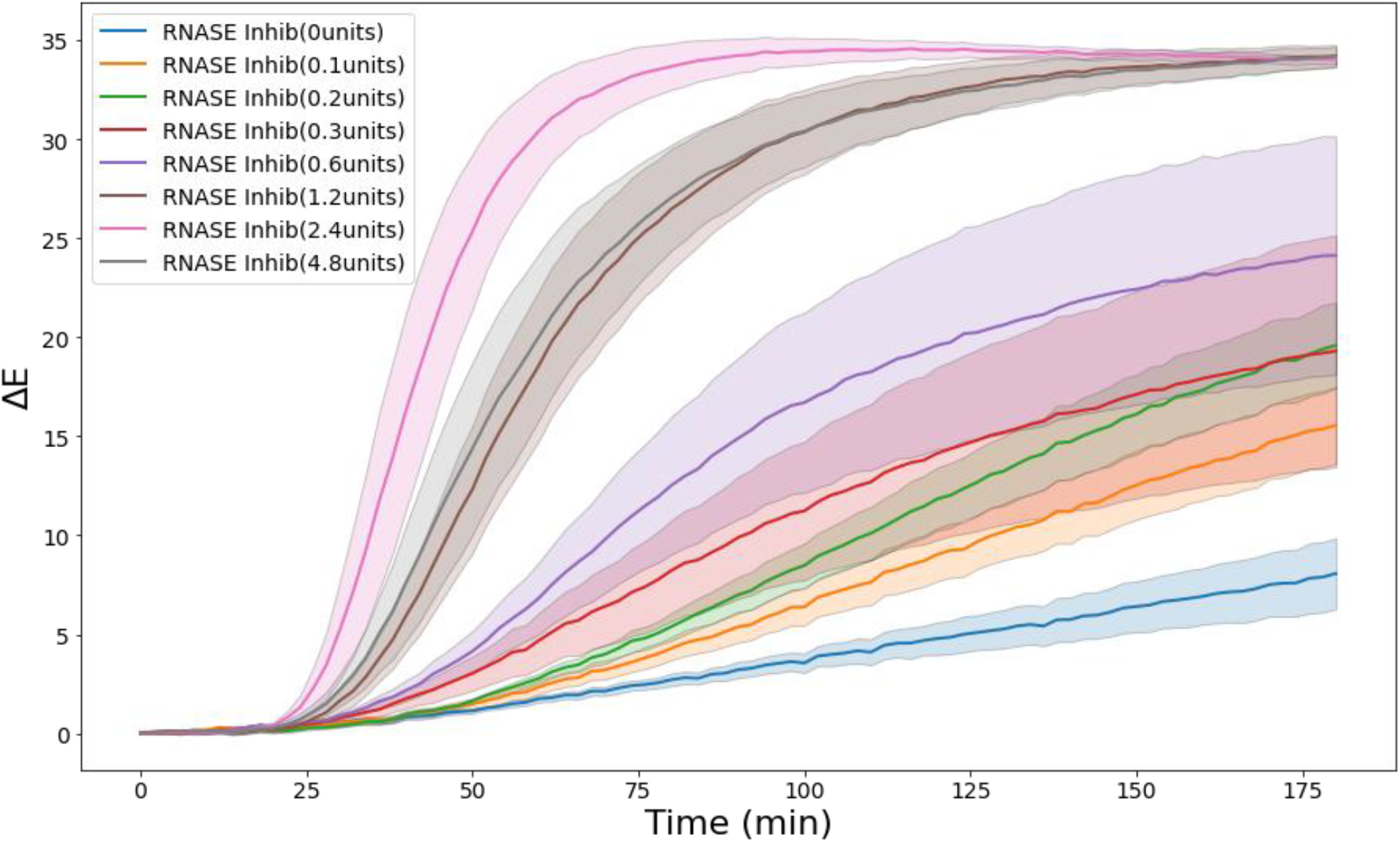
Effect of increasing amounts of RNASE Inhibitor added to the reaction mixture.

**Figure S5:**
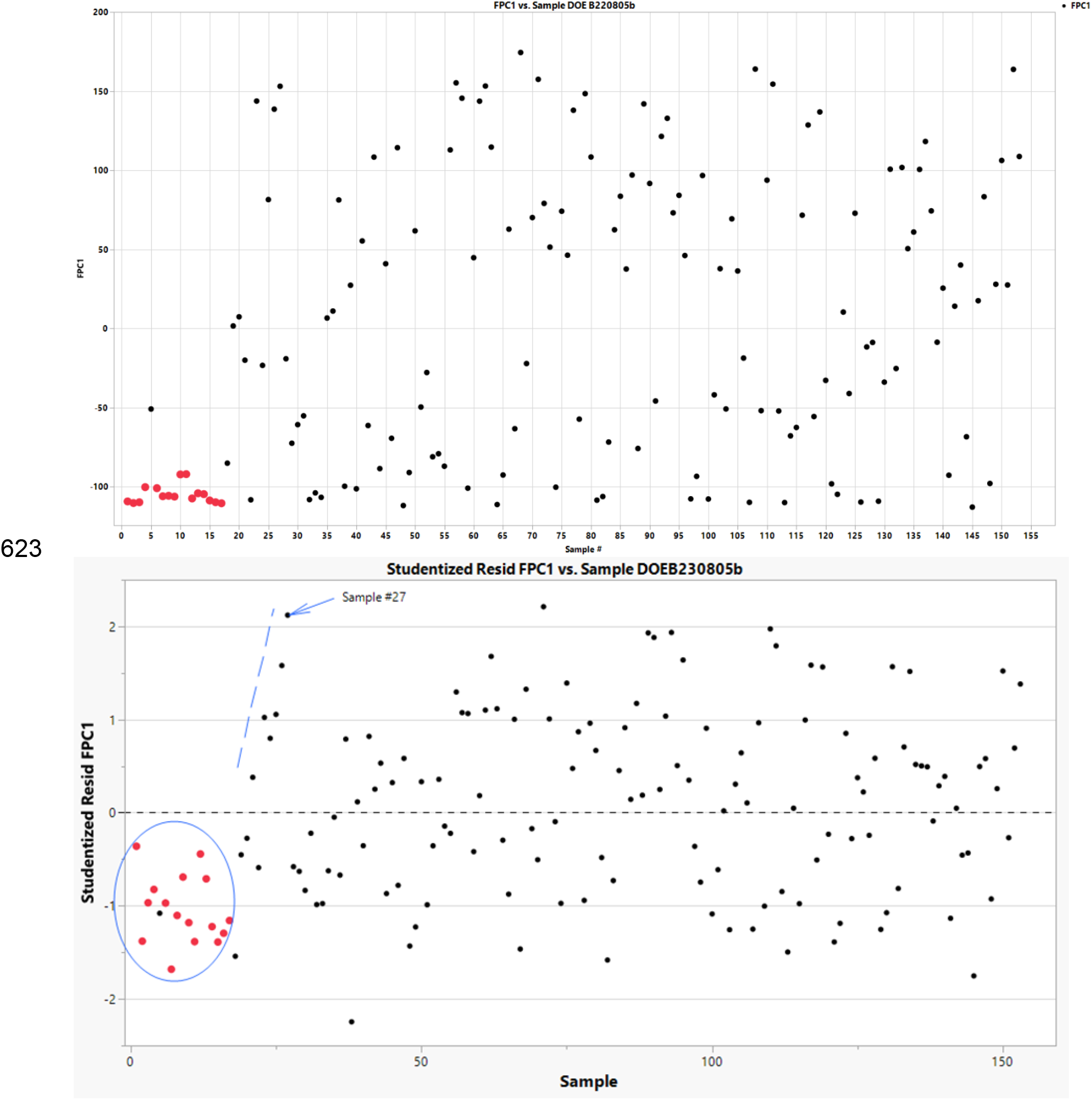
Extracted FPC1 values as a function of DOE number from PANOx-SP components added to baseline Cai. (top) Raw FPC1 values. (bottom) Studentized residuals. Red dots indicate region of apparent of unexpected systematic bias.

**Figure S6:**
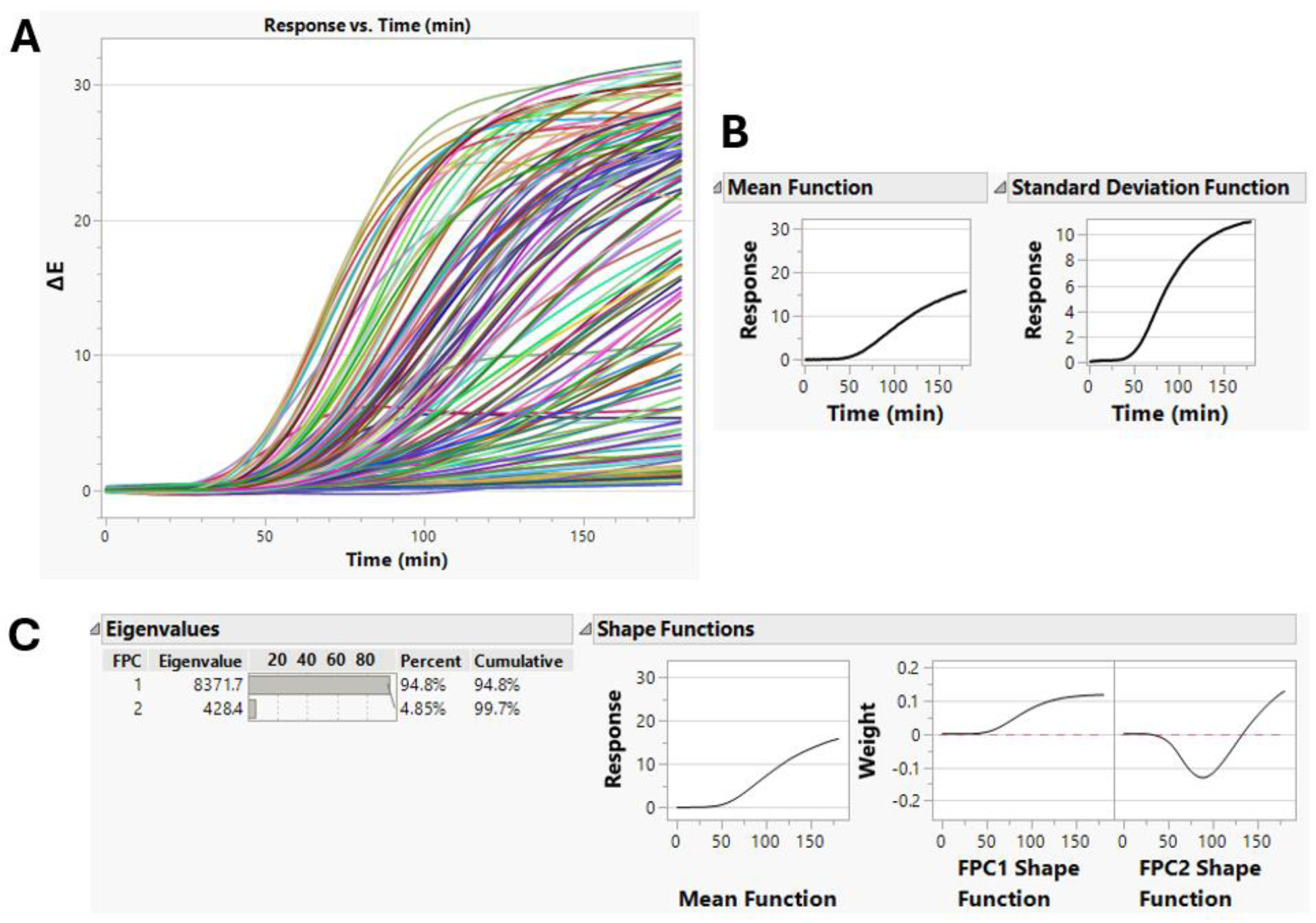
Functional Principal Component decomposition of curves resulting from DOE of adding PANOx-SP components to baseline Cai reactions.(A) Raw traces of ΔE over time extracted from images. (B) Mean and standard deviation functions of the full data set after removal of outliers. (C) Results of FPC analysis.

**Figure S7:**
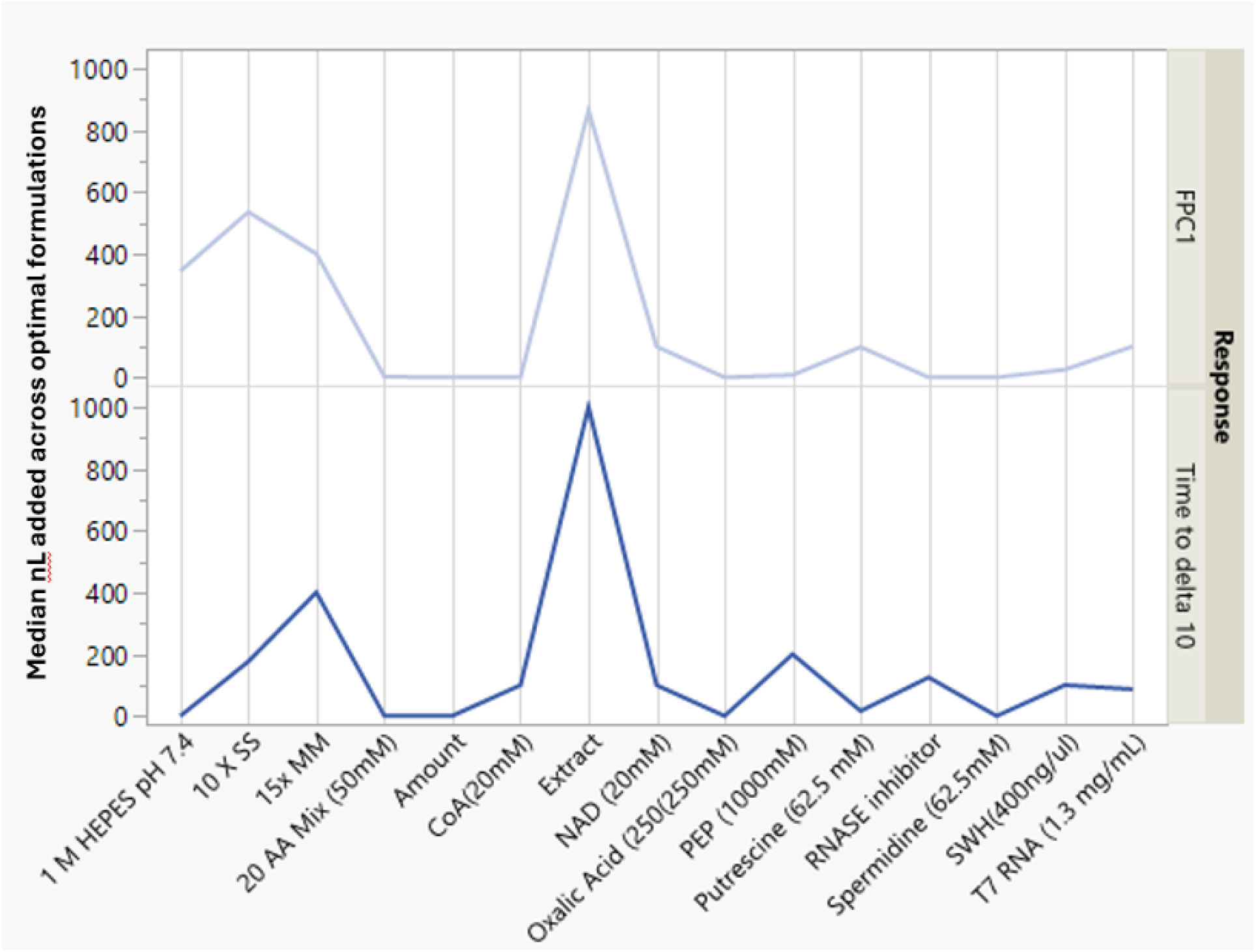
Median volumes of each component added in the predicted optimal formulations based on optimizing for FPC1 (top) or time to ΔE≥10 (bottom).

**Figure S8.**
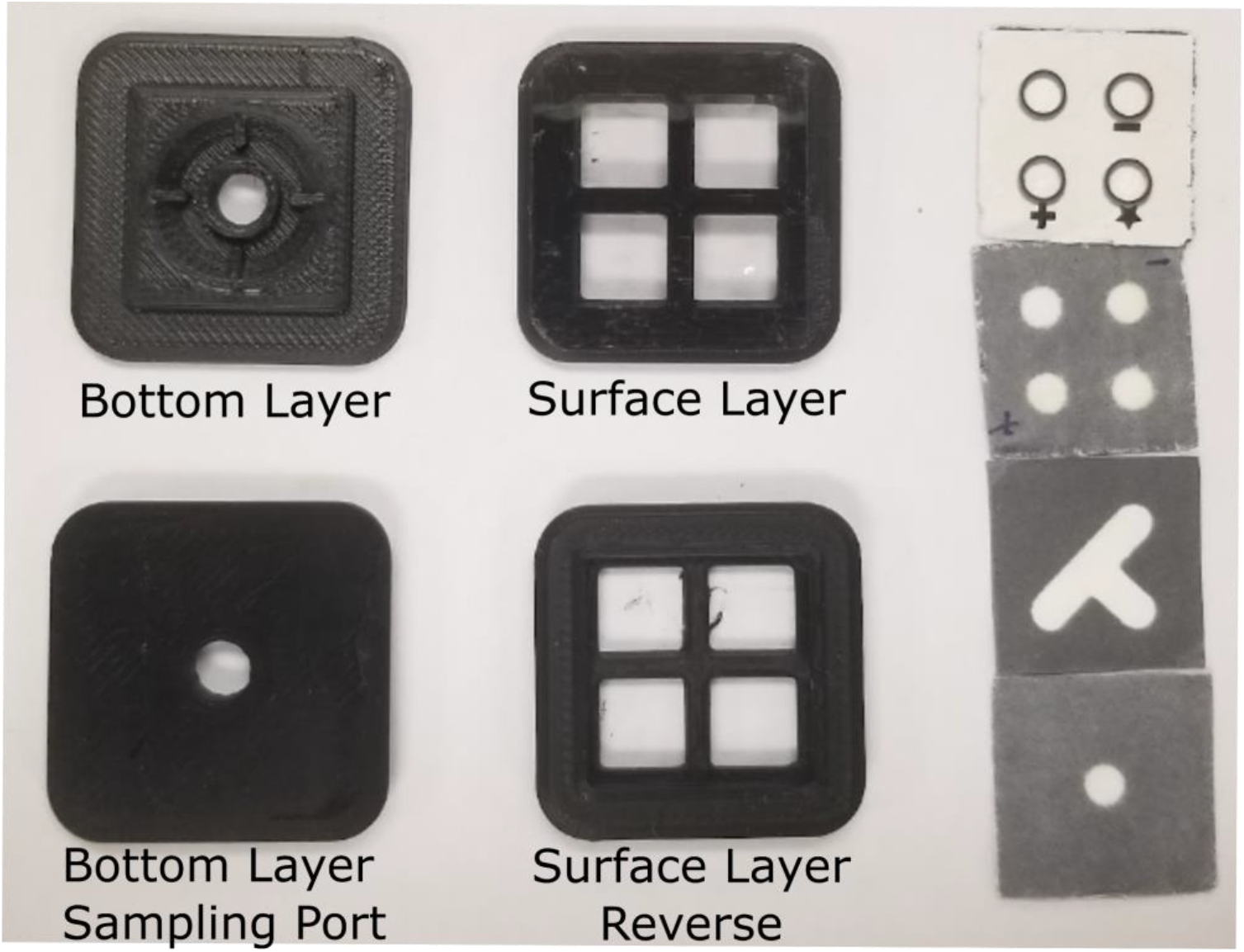
Fitted bottom and surface layers hold together the paper fluidics. Sample is added directly to the sampling port and visualized on the surface layer covered in transparent film.

**Table S1:** Cost comparison of CFE reactions using Cai and PANOx-SP formulations. Media calculations assume 300 mL per L, and 3 mL of lysate made per L of media.

|  | Cai | PANox-SP |
| --- | --- | --- |
| *Media Components for Lysate* |  |  |
| Tryptone | \$ 77.10 | \$ 77.10 |
| Yeast Extract | \$ 34.80 | \$ 34.80 |
| NaCl | \$ 3.41 | \$ 3.41 |
| Potassium phosphate, monobasic | \$ 10.75 | \$ 10.75 |
| Potassium phosphate, monobasic | \$ 10.26 | \$ 10.26 |
| DTT | \$ 87.41 | \$ 87.41 |
| *CFE Components* |  |  |
| YNB | \$ - | \$ - |
| HEPES | \$ - | \$ 5.79 |
| Tris | \$ - | \$ - |
| Potassium Phosphate | \$ 0.16 | \$ - |
| ATP | \$ - | \$ 40.67 |
| GTP | \$ - | \$ 302.40 |
| CTP | \$ - | \$ 227.61 |
| UTP | \$ - | \$ 469.13 |
| AMP | \$ 6.52 | \$ - |
| GMP | \$ 3.54 | \$ - |
| CMP | \$ 17.57 | \$ - |
| UMP | \$ 5.57 | \$ - |
| Oxidized glutathione | \$ 5.69 | \$ - |
| Tyrosine | \$ 0.12 | \$ - |
| PEP | \$ - | \$2,142.72 |
| NAD | \$ - | \$ 728.32 |
| CoA | \$ - | \$ 681.80 |
| Potassium Glutamate | \$ 16.01 | \$ 10.78 |
| Ammonium Glutamate | \$ - | \$ 0.62 |
| Magnesium Glutamate | \$ 8.62 | \$ 17.25 |
| Sodium/Potassium Oxalate | \$ 0.05 | \$ 0.05 |
| Putrescine | \$ - | \$ 0.70 |
| Spermidine | \$ 5.05 | \$ 5.05 |
| Alanine | \$ 0.11 | \$ 0.11 |
| Arginine | \$ 0.13 | \$ 0.13 |
| Asparagine | \$ 0.20 | \$ 0.20 |
| Aspartic acid | \$ 0.03 | \$ 0.03 |
| Cysteine | \$ 0.19 | \$ 0.19 |
| Glutamic acid | \$ 0.02 | \$ 0.02 |
| Glutamine | \$ 0.16 | \$ 0.16 |
| Glycine | \$ 0.02 | \$ 0.02 |
| Histidine | \$ 0.17 | \$ 0.17 |
| Isoleucine | \$ 0.25 | \$ 0.25 |
| Leucine | \$ 0.15 | \$ 0.15 |
| Lysine | \$ 0.99 | \$ 0.99 |
| Methionine | \$ 0.11 | \$ 0.11 |
| Phenylalanine | \$ 0.19 | \$ 0.19 |
| Proline | \$ 0.13 | \$ 0.13 |
| Serine | \$ 0.15 | \$ 0.15 |
| Threonine | \$ 0.25 | \$ 0.25 |
| Tryptophan | \$ 0.31 | \$ 0.31 |
| Tyrosine | \$ 0.26 | \$ 0.26 |
| Valine | \$ 0.12 | \$ 0.12 |
| Folinic Acid | \$ - | \$ 89.25 |
| E. coli tRNA | \$ - | \$ 246.58 |
| DNA Template | \$ 92.03 | \$ 92.03 |
| T7 RNA polymerase | \$ 56.40 | \$ 282.00 |
| **Total:** | **\$444.97** | **\$5,570.37** |

**Table S2:** Energy mixture components for PANOx-SP base reactions.

| **Component** | **PANOx-SP Final Concentration** |
| --- | --- |
| Magnesium | 12 mM |
| Potassium glutamate | 130 mM |
| Ammonium glutamate | 10 mM |
| Magnesium acetate | 3.7 mM |
| Potassium acetate | 6.2 mM |
| Tris acetate | 5.67 mM (pH 8.2) |
| HEPES | 7.5 mM (pH 7.4) |
| ATP | 1.2 mM |
| GTP | 0.85 mM |
| CTP | 0.85 mM |
| UTP | 0.85 mM |
| Folic acid | 0.072 mM |
| tRNA | 170.6 µg/mL |
| Alanine | 2 mM |
| Arginine | 2 mM |
| Histidine | 2 mM |
| Lysine (monoHCl) | 2 mM |
| Aspartic acid | 2 mM |
| Glutamic acid | 2 mM |
| Isoleucine | 2 mM |
| Leucine | 2 mM |
| Methionine | 2 mM |
| Phenylalanine | 2 mM |
| Tryptophan | 2 mM |
| Tyrosine | 2 mM |
| Valine | 2 mM |
| Serine | 2 mM |
| Threonine | 2 mM |
| Asparagine | 2 mM |
| Glutamine | 2 mM |
| Cysteine | 2 mM |
| Glycine | 2 mM |
| Proline | 2 mM |
| PEP | 33 mM |
| NAD | 0.33 mM |
| CoA | 0.27 mM |
| Spermidine | 1.5 mM |
| Putrescine | 2 mM |
| Oxalic acid | 4 mM |
| T7 RNA polymerase | 100 µg/mL |
| Plasmid DNA | 6.4 nM |
| RNase Inhibitor | 1.2 U/µL |
| Cell extract | 30% v/v |
| DTT | 0.5 mM |

**Table S3.** Energy mixture components for Cai base reactions.

| **Component** | **Concentration** |
| --- | --- |
| AMP | 1.2 mM |
| UMP | 0.86 mM |
| CMP | 0.86 mM |
| GMP | 0.86 mM |
| T7 RNA Polymerase | 0.02 mg/mL |
| Magnesium Glutamate | 8 mM |
| Potassium Glutamate | 260 mM |
| Potassium Oxalate | 4 mM |
| Potassium Phosphate pH 7.0 | 15 mM |
| Spermidine | 1.5 mM |
| Oxidized Glutathione | 2 mM |
| Alanine | 2 mM |
| Arginine | 2 mM |
| Histidine | 2 mM |
| Lysine (monoHCl) | 2 mM |
| Aspartic acid | 2 mM |
| Glutamic acid | 2 mM |
| Isoleucine | 2 mM |
| Leucine | 2 mM |
| Methionine | 2 mM |
| Phenylalanine | 2 mM |
| Tryptophan | 1 mM |
| Tyrosine | 2 mM |
| Valine | 2 mM |
| Serine | 2 mM |
| Threonine | 2 mM |
| Asparagine | 2 mM |
| Glutamine | 2 mM |
| Cysteine | 2 mM |
| Glycine | 2 mM |
| Proline | 2 mM |
| Cell extract | 30% v/v |
| RNase Inhibitor | 1.2 U/μL |

**Table S4.** Working solutions used in energy mixture components for PANOx-SP reagent additives.

| Reagent | Concentration |
| --- | --- |
| Salt Solution (SS) | 10X |
| HEPES | 1 M |
| Cell-Free Master Mix (MM) | 15X |
| AA Mix | 50 mM |
| PEP | 1 M |
| NAD | 20 mM |
| CoA | 20 mM |
| Oxalic Acid | 250 mM |
| Putrescine | 62.5 mM |
| Spermidine | 62.5 mM |
| T7 RNAP | 1.3 mg/mL |
| RNASE inhibitor Murine | 4 units/μL |
| Extract | ~30 mg/mL |
| CPRG | - |
| DNA (each plasmid) | 400 ng/μl |

**Table S5:** Working solutions used in energy mixture components for Cai reagent additives.

| Reagent | Concentration |
| --- | --- |
| L-Glutamic acid hemimagnesium salt tetrahydrate | 40 mM |
| L-Glutamic acid monopotassium salt monohydrate | 1300 mM |
| AMP | 25 mM |
| GMP | 25 mM |
| UMP | 25 mM |
| CMP | 25 mM |
| Oxalic Acid | 150 mM |
| L-Glutathione oxidized | 50 mM |
| Spermidine | 125 mM |
| Potassium phosphate ($K_2HPO_4$) | 307.5 mM |
| Potassium phosphate (KH<sub>2</sub>PO<sub>4</sub>) | 192.5 mM |
| Amino Acids (19) | 25 mM |
| Tyrosine | 25 mM |
| T7 RNAP | 1.3 mg/mL |
| RNASE inhibitor Murine | - |
| Lysate | ~30 mg/mL |
| CPRG | - |
| DNA (each plasmid) | 400 ng/μl |

**Table S6:** Excipients tested for heat tolerance.

| Excipient | Concentration | Role | Reference |
| --- | --- | --- | --- |
| Dextran 70 | 50 mg/mL | Cryoprotectant | *Hagen et al. (1997)* |
| PEG8K | 5% | Lyoprotectant | *Manning et al. (2010)* |
| Maltose | 50 mM | Cryoprotectant | *Chirife et al. (2000)* |
| H<sub>2</sub>KPO<sub>4</sub> | 50 mM | Buffering agent | *Pikal-Cleland et al. (2000)* |
| Ficoll | 50 mg/mL | Cryoprotectant | *Wang et al. (2006)* |
| Maltodextrin | 50 mM | Bulking agent | *Patel and Pikal (1996)* |
| Raffinose | 50 mM | Cryoprotectant | *Wang et al. (2006)* |
| Lactose | 50 mM | Cryoprotectant | *Wang et al. (2006)* |
| Rhamnose | 50 mM | Cryoprotectant | *Chirife et al. (2000)* |
| Mannitol | 50 mM | Cryoprotectant | *Wang et al. (2006)* |
| Sucrose | 50 mM | Cryoprotectant | *Wang et al. (2006)* |
| Malitol | 50 mM | Cryoprotectant | *Wang et al. (2006)* |
| Sorbitol | 50 mM | Cryoprotectant | *Chirife et al. (2000)* |
| TMAO | 50 mM | Stabilizer/Protectant | *Wüstner and Solanko (2015)* |

**Table S7:** Linear blending coefficients from the DOE model of PANOx-SP components added to baseline Cai. Positive coefficients indicate an increase in FPC1 as the proportion of the component changes.

| Linear Blending Coefficient | Name |
| --- | --- |
| 133.6 | 1 M HEPES pH 7.4 |
| 124.7 | Oxalic Acid (250(250mM) |
| 117.1 | SWH(400ng/μl) |
| 88.7 | T7 RNA (1.3 mg/mL) |
| 70.1 | CoA(20mM) |
| 61.2 | 15x MM |
| 44.6 | Putrescine (62.5 mM) |
| 23.3 | NAD (20mM) |
| 10.9 | Extract |
| -2.7 | Spermidine (62.5mM) |
| -21.8 | 10 X SS |
| -23.9 | 20 AA Mix (50mM) |
| -115.2 | RNASE inhibitor |
| -151.5 | PEP (1000mM) |

**Table S8:** Non-linear blending coefficients from the DOE model of PANOx-SP components added to baseline Cai. Positive coefficients indicate an increase in FPC1 as the proportion of the component changes.

| Non-linear Blending Coefficient | Component Names |
| --- | --- |
| 1,654.40 | Spermidine (62.5mM)*RNASE inhibitor |
| 1,173.80 | CoA(20mM)*RNASE inhibitor |
| 1,027.40 | PEP (1000mM)*RNASE inhibitor |
| 901.2 | 20 AA Mix (50mM)*Spermidine (62.5mM) |
| 522.3 | 10 X SS*SWH(400ng/μl) |
| 330.2 | PEP (1000mM) * Extract |
| 242 | 10 X SS*PEP (1000mM) |
| 17.9 | PEP (1000mM) * amount (total added) |
| 6.7 | Extract *amount (total added) |
| -45.1 | SWH(400ng/μl)*amount (total added) |
| -72.8 | Putrescine (62.5 mM) * amount (total added) |
| -196.2 | 1 M HEPES pH 7.4*Extract |
| -383 | 1 M HEPES pH 7.4 *PEP (1000mM) |
| -628.6 | 15x MM *CoA(20mM) |
| -967.3 | 1 M HEPES pH 7.4*Spermidine (62.5mM) |
| -1,361.90 | 15x MM*Oxalic Acid (250(250mM) |
| -1,541.70 | CoA(20mM)*Oxalic Acid (250(250mM) |
| -1,565.00 | 1 M HEPES pH 7.4 * SWH(400ng/μl) |
| -2,299.30 | Putrescine (62.5 mM) * SWH(400ng/μl) |
| -2,708.30 | Spermidine (62.5mM)* T7 RNA (1.3 mg/mL) |

**Table S9:** Results of independent t-tests performed to compare the PANOx-SP controls against each experimental dataset when ΔE≥10 for the optimized Cai formulations. The p-values assess the statistical significance of these differences using n=4 samples.

| Comparison | p-value |
| --- | --- |
| Cai+Opt1 | 0.415247 |
| Cai+Opt2 | 0.650334 |
| Cai+Opt3 | 0.256598 |
| Cai+Opt4 | 0.816178 |
| Cai+Opt5 | 0.871965 |
| Cai+Opt6 | 0.96559 |
| Cai+Opt7 | 0.252361 |
| Cai+Opt8 | 0.407565 |
| Cai+Opt9 | 0.735271 |

**Table S10:** Cost breakdown of Cai optimized formulations. Costs are given per 1 µL reaction.

| Index | Cai-Opt Costs | Baseline Cai Cost | Total Cost per Reaction |
| --- | --- | --- | --- |
| 1 | \$ 1.95876 | \$ 0.00044 | \$ 1.95920 |
| 2 | \$ 0.00129 | \$ 0.00044 | \$ 0.00173 |
| 3 | \$ 0.00133 | \$ 0.00044 | \$ 0.00177 |
| 4 | \$ 1.96275 | \$ 0.00044 | \$ 1.96320 |
| 5 | \$ 0.00132 | \$ 0.00044 | \$ 0.00177 |
| 6 | \$ 0.00131 | \$ 0.00044 | \$ 0.00176 |
| 7 | \$ 0.49299 | \$ 0.00044 | \$ 0.49343 |
| 8 | \$ 1.96420 | \$ 0.00044 | \$ 1.96465 |
| 9 | \$ 0.00141 | \$ 0.00044 | \$ 0.00186 |
| 10 | \$ 1.95528 | \$ 0.00044 | \$ 1.95573 |
| 11 | \$ 0.00021 | \$ 0.00044 | \$ 0.00065 |
| 12 | \$ 0.49558 | \$ 0.00044 | \$ 0.49603 |
| 13 | \$ 0.49721 | \$ 0.00044 | \$ 0.49766 |
| 14 | \$ 0.00022 | \$ 0.00044 | \$ 0.00067 |
| 15 | \$ 1.95924 | \$ 0.00044 | \$ 1.95968 |
| 16 | \$ 0.00259 | \$ 0.00044 | \$ 0.00303 |
| 17 | \$ 1.95974 | \$ 0.00044 | \$ 1.96018 |
| 18 | \$ 0.00255 | \$ 0.00044 | \$ 0.00300 |
| 19 | \$ 0.49694 | \$ 0.00044 | \$ 0.49738 |
| 20 | \$ 0.50248 | \$ 0.00044 | \$ 0.50293 |
| 21 | \$ 0.49139 | \$ 0.00044 | \$ 0.49184 |
| 22 | \$ 1.96049 | \$ 0.00044 | \$ 1.96094 |
| 23 | \$ 1.95581 | \$ 0.00044 | \$ 1.95625 |
| 24 | \$ 1.96259 | \$ 0.00044 | \$ 1.96304 |
| 25 | \$ 1.47543 | \$ 0.00044 | \$ 1.47588 |
| 26 | \$ 0.00316 | \$ 0.00044 | \$ 0.00361 |
| 27 | \$ 1.96094 | \$ 0.00044 | \$ 1.96138 |
| 28 | \$ 0.00442 | \$ 0.00044 | \$ 0.00487 |
| 29 | \$ 0.97770 | \$ 0.00044 | \$ 0.97814 |
| 30 | \$ 1.96160 | \$ 0.00044 | \$ 1.96204 |

**Table S11:** Linear blending coefficients from the DOE model of heat tolerance excipients added to baseline PANOx-SP. Positive coefficients indicate an increase in FPC1 as the proportion of the component changes.

| Linear Blending Coefficient | Component Name |
| --- | --- |
| 258.4 | Raffinose 50mM |
| 170.5 | Maltose 50mM |
| 136.9 | Ficoll 50mM |
| 132.4 | TMAO 50mM |
| 131.2 | Maltodextrin 50mM |
| 72.3 | Sorbitol 50mM |
| 33.5 | H2KPO4 |
| 18.8 | Dextran 70 50 mM |
| -62.6 | Sucrose 50mM |
| -107.8 | Mannitol 50mM |
| -126.1 | Lactose 50mM |
| -135.7 | Malitol 50mM |
| -248.4 | PEG5% |
| -290.8 | Rhamnose 50mM |

**Table S12:** Non-linear blending coefficients from the DOE model of heat tolerance excipients added to baseline PANOx-SP. Positive coefficients indicate an increase in FPC1 as the proportion of the component changes.

| Non-linear Blending Coefficient | Component Names |
| --- | --- |
| 1580.7 | Dextran 70 50 mM*Mannitol 50mM |
| 1453.4 | PEG5%*Rhamnose 50mM |
| 1203.2 | Sucrose 50mM*Malitol 50mM |
| 1158.8 | Lactose 50mM*Rhamnose 50mM |
| 1136.7 | Maltodextrin 50mM*Sucrose 50mM |
| 1119.7 | H2KPO4*Ficoll 50mM |
| 996.2 | Mannitol 50mM*Malitol 50mM |
| 751.6 | H2KPO4*Raffinose 50mM |
| 669.2 | Ficoll 50mM*Rhamnose 50mM |
| 389.6 | Maltose 50mM*Malitol 50mM |
| -443.1 | Dextran 70 50 mM*Malitol 50mM |
| -659.5 | PEG5%*Sucrose 50mM |
| -729.5 | Dextran 70 50 mM*Ficoll 50mM |
| -735.3 | Maltose 50mM*Ficoll 50mM |
| -750.8 | Maltose 50mM*H2KPO4 |
| -853.5 | Maltose 50mM*Maltodextrin 50mM |
| -1038.9 | Raffinose 50mM*Sorbitol 50mM |
| -1100.2 | Maltodextrin 50mM*TMAO 50mM |
| -1120.1 | Maltose 50mM*TMAO 50mM |
| -1423.1 | Ficoll 50mM*Rhamnose 50mM |

**Table S13.** Results of independent t-tests performed to compare the point where of ΔE≥10 for PANOx-SP dataset from the Cai optimization experiment against the heat optimized formulations (Heat-Opt1 through Heat-Opt14). The p-values assess the statistical significance of these differences using n=4 samples.

| Comparison | p-value |
| --- | --- |
| PANOx-SP vs Heat-Opt1 | 0.94 |
| PANOx -SP vs Heat-Opt2 | 0.99 |
| PANOx -SP vs Heat-Opt3 | 0.96 |
| PANOx -SP vs Heat-Opt4 | 0.98 |
| PANOx -SP vs Heat-Opt5 | 0.83 |
| PANOx -SP vs Heat-Opt6 | 0.96 |
| PANOx -SP vs Heat-Opt7 | 0.99 |
| PANOx -SP vs Heat-Opt8 | 0.97 |
| PANOx -SP vs Heat-Opt9 | 0.79 |
| PANOx -SP vs Heat-Opt10 | 0.97 |
| PANOx -SP vs Heat-Opt11 | 0.97 |
| PANOx -SP vs Heat-Opt12 | 0.97 |
| PANOx -SP vs Heat-Opt13 | 0.92 |
| PANOx -SP vs Heat-Opt14 | 0.87 |

## References

(1) Lavickova, B.; Maerkl, S. J. A Simple, Robust, and Low-Cost Method To Produce the PURE Cell-Free System. ACS Synth. Biol. 2019, 8 (2), 455–462. 10.1021/acssynbio.8b00427.

(2) Dopp, B. J. L.; Tamiev, D. D.; Reuel, N. F. Cell-Free Supplement Mixtures: Elucidating the History and Biochemical Utility of Additives Used to Support in Vitro Protein Synthesis in E. Coli Extract. Biotechnology Advances 2019, 37 (1), 246–258. 10.1016/j.biotechadv.2018.12.006.

(3) Amalfitano, E.; Karlikow, M.; Norouzi, M.; Jaenes, K.; Cicek, S.; Masum, F.; Sadat Mousavi, P.; Guo, Y.; Tang, L.; Sydor, A.; Ma, D.; Pearson, J. D.; Trcka, D.; Pinette, M.; Ambagala, A.; Babiuk, S.; Pickering, B.; Wrana, J.; Bremner, R.; Mazzulli, T.; Sinton, D.; Brumell, J. H.; Green, A. A.; Pardee, K. A Glucose Meter Interface for Point-of-Care Gene Circuit-Based Diagnostics. Nat Commun 2021, 12 (1), 724. 10.1038/s41467-020-20639-6.

(4) Pardee, K.; Green, A. A.; Ferrante, T.; Cameron, D. E.; Daleykeyser, A.; Yin, P.; Collins, J. J. Paper-Based Synthetic Gene Networks. Cell 2014, 159 (4), 940–954. 10.1016/j.cell.2014.10.004.

(5) Beabout, K.; Bernhards, C. B.; Thakur, M.; Turner, K. B.; Cole, S. D.; Walper, S. A.; Chávez, J. L.; Lux, M. W. Optimization of Heavy Metal Sensors Based on Transcription Factors and Cell-Free Expression Systems. ACS Synth. Biol. 2021, 10 (11), 3040–3054. 10.1021/acssynbio.1c00331.

(6) McNerney, M. P.; Zhang, Y.; Steppe, P.; Silverman, A. D.; Jewett, M. C.; Styczynski, M. P. Point-of-Care Biomarker Quantification Enabled by Sample-Specific Calibration. Sci. Adv. 2019, 5 (9), eaax4473. 10.1126/sciadv.aax4473.

(7) Sadat Mousavi, P.; Smith, S. J.; Chen, J. B.; Karlikow, M.; Tinafar, A.; Robinson, C.; Liu, W.; Ma, D.; Green, A. A.; Kelley, S. O.; Pardee, K. A Multiplexed, Electrochemical Interface for Gene-Circuit-Based Sensors. Nat. Chem. 2020, 12 (1), 48–55. 10.1038/s41557-019-0366-y.

(8) Pandi, A.; Grigoras, I.; Borkowski, O.; Faulon, J.-L. Optimizing Cell-Free Biosensors to Monitor Enzymatic Production. ACS Synth. Biol. 2019, 8 (8), 1952–1957. 10.1021/acssynbio.9b00160.

(9) Whitfield, C. J.; Banks, A. M.; Dura, G.; Love, J.; Fieldsend, J. E.; Goodchild, S. A.; Fulton, D. A.; Howard, T. P. Cell-Free Protein Synthesis in Hydrogel Materials. Chem. Commun. 2020, 56 (52), 7108–7111. 10.1039/D0CC02582H.

(10) Hunt, J. P.; Zhao, E. L.; Free, T. J.; Soltani, M.; Warr, C. A.; Benedict, A. B.; Takahashi, M. K.; Griffitts, J. S.; Pitt, W. G.; Bundy, B. C. Towards Detection of SARS-CoV-2 RNA in Human Saliva: A Paper-Based Cell-Free Toehold Switch Biosensor with a Visual Bioluminescent Output. New Biotechnology 2022, 66, 53–60. 10.1016/j.nbt.2021.09.002.

(11) Pardee, K. Perspective: Solidifying the Impact of Cell-Free Synthetic Biology through Lyophilization. Biochemical Engineering Journal 2018, 138, 91–97. 10.1016/j.bej.2018.07.008.

(12) Sharpes, C. E.; McManus, J. B.; Blum, S. M.; Mgboji, G. E.; Lux, M. W. Assessment of Colorimetric Reporter Enzymes in the PURE System. ACS Synth. Biol. 2021, 10 (11), 3205–3208. 10.1021/acssynbio.1c00360.

(13) Arce, A.; Guzman Chavez, F.; Gandini, C.; Puig, J.; Matute, T.; Haseloff, J.; Dalchau, N.; Molloy, J.; Pardee, K.; Federici, F. Decentralizing Cell-Free RNA Sensing With the Use of Low-Cost Cell Extracts. Front. Bioeng. Biotechnol. 2021, 9, 727584. 10.3389/fbioe.2021.727584.

(14) Karig, D. K.; Bessling, S.; Thielen, P.; Zhang, S.; Wolfe, J. Preservation of Protein Expression Systems at Elevated Temperatures for Portable Therapeutic Production. J. R. Soc. Interface. 2017, 14 (129), 20161039. 10.1098/rsif.2016.1039.

(15) Gregorio, N. E.; Kao, W. Y.; Williams, L. C.; Hight, C. M.; Patel, P.; Watts, K. R.; Oza, J. P. Unlocking Applications of Cell-Free Biotechnology through Enhanced Shelf Life and Productivity of *E. Coli* Extracts. ACS Synth. Biol. 2020, 9 (4), 766–778. 10.1021/acssynbio.9b00433.

(16) Smith, M. T.; Berkheimer, S. D.; Werner, C. J.; Bundy, B. C. Lyophilized *Escherichia Coli* - Based Cell-Free Systems for Robust, High-Density, Long-Term Storage. BioTechniques 2014, 56 (4), 186–193. 10.2144/000114158.

(17) Cai, Q.; Hanson, J. A.; Steiner, A. R.; Tran, C.; Masikat, M. R.; Chen, R.; Zawada, J. F.; Sato, A. K.; Hallam, T. J.; Yin, G. A Simplified and Robust Protocol for Immunoglobulin Expression in *E Scherichia Coli* Cell-free Protein Synthesis Systems. Biotechnology Progress 2015, 31 (3), 823–831. 10.1002/btpr.2082.

(18) Jeong, S.; González-Grandío, E.; Navarro, N.; Pinals, R. L.; Ledesma, F.; Yang, D.; Landry, M. P. Extraction of Viral Nucleic Acids with Carbon Nanotubes Increases SARS-CoV-2 Quantitative Reverse Transcription Polymerase Chain Reaction Detection Sensitivity. ACS Nano 2021, 15 (6), 10309–10317. 10.1021/acsnano.1c02494.

(19) Brouillette, M. Repurposing the Cell Engine: Northwestern Announces the Cell-Free Biomanufacturing Institute. GEN Biotechnology 2022, 1 (3), 218–221. 10.1089/genbio.2022.29037.mbr.

(20) Borkowski, O.; Koch, M.; Zettor, A.; Pandi, A.; Batista, A. C.; Soudier, P.; Faulon, J.-L. Large Scale Active-Learning-Guided Exploration for in Vitro Protein Production Optimization. Nat Commun 2020, 11 (1), 1872. 10.1038/s41467-020-15798-5.

(21) Moore, S. J.; MacDonald, J. T.; Wienecke, S.; Ishwarbhai, A.; Tsipa, A.; Aw, R.; Kylilis, N.; Bell, D. J.; McClymont, D. W.; Jensen, K.; Polizzi, K. M.; Biedendieck, R.; Freemont, P. S. Rapid Acquisition and Model-Based Analysis of Cell-Free Transcription–Translation Reactions from Nonmodel Bacteria. Proc. Natl. Acad. Sci. U.S.A. 2018, 115 (19). 10.1073/pnas.1715806115.

(22) Karim, A. S.; Dudley, Q. M.; Juminaga, A.; Yuan, Y.; Crowe, S. A.; Heggestad, J. T.; Garg, S.; Abdalla, T.; Grubbe, W. S.; Rasor, B. J.; Coar, D. N.; Torculas, M.; Krein, M.; Liew, F.; Quattlebaum, A.; Jensen, R. O.; Stuart, J. A.; Simpson, S. D.; Köpke, M.; Jewett, M. C. In Vitro Prototyping and Rapid Optimization of Biosynthetic Enzymes for Cell Design. Nat Chem Biol 2020, 16 (8), 912–919. 10.1038/s41589-020-0559-0.

(23) Banks, A. M.; Whitfield, C. J.; Brown, S. R.; Fulton, D. A.; Goodchild, S. A.; Grant, C.; Love, J.; Lendrem, D. W.; Fieldsend, J. E.; Howard, T. P. Key Reaction Components Affect the Kinetics and Performance Robustness of Cell-Free Protein Synthesis Reactions. Computational and Structural Biotechnology Journal 2022, 20, 218–229. 10.1016/j.csbj.2021.12.013.

(24) Gilman, J.; Walls, L.; Bandiera, L.; Menolascina, F. Statistical Design of Experiments for Synthetic Biology. ACS Synth. Biol. 2021, 10 (1), 1–18. 10.1021/acssynbio.0c00385.

(25) Malcı, K.; Walls, L. E.; Rios-Solis, L. Rational Design of CRISPR/Cas12a-RPA Based One-Pot COVID-19 Detection with Design of Experiments. ACS Synth. Biol. 2022, 11 (4), 1555– 1567. 10.1021/acssynbio.1c00617.

(26) Pardee, K.; Green, A. A.; Takahashi, M. K.; Braff, D.; Lambert, G.; Lee, J. W.; Ferrante, T.; Ma, D.; Donghia, N.; Fan, M.; Daringer, N. M.; Bosch, I.; Dudley, D. M.; O’Connor, D. H.; Gehrke, L.; Collins, J. J. Rapid, Low-Cost Detection of Zika Virus Using Programmable Biomolecular Components. Cell 2016, 165 (5), 1255–1266. 10.1016/j.cell.2016.04.059.

(27) Blum, S. M.; Lee, M. S.; Mgboji, G. E.; Funk, V. L.; Beabout, K.; Harbaugh, S. V.; Roth, P. A.; Liem, A. T.; Miklos, A. E.; Emanuel, P. A.; Walper, S. A.; Chávez, J. L.; Lux, M. W. Impact of Porous Matrices and Concentration by Lyophilization on Cell-Free Expression. ACS Synth. Biol. 2021, 10 (5), 1116–1131. 10.1021/acssynbio.0c00634.

(28) Lopreside, A.; Wan, X.; Michelini, E.; Roda, A.; Wang, B. Comprehensive Profiling of Diverse Genetic Reporters with Application to Whole-Cell and Cell-Free Biosensors. Anal. Chem. 2019, 91 (23), 15284–15292. 10.1021/acs.analchem.9b04444.

(29) Siegal-Gaskins, D.; Tuza, Z. A.; Kim, J.; Noireaux, V.; Murray, R. M. Gene Circuit Performance Characterization and Resource Usage in a Cell-Free “Breadboard.” ACS Synth. Biol. 2014, 3 (6), 416–425. 10.1021/sb400203p.

(30) Haeuser, C.; Goldbach, P.; Huwyler, J.; Friess, W.; Allmendinger, A. Impact of Dextran on Thermal Properties, Product Quality Attributes, and Monoclonal Antibody Stability in Freeze-Dried Formulations. European Journal of Pharmaceutics and Biopharmaceutics 2020, 147, 45–56. 10.1016/j.ejpb.2019.12.010.

(31) Johnson, R. E.; Kirchhoff, C. F.; Gaud, H. T. Mannitol–Sucrose Mixtures—Versatile Formulations for Protein Lyophilization. Journal of Pharmaceutical Sciences 2002, 91 (4), 914–922. 10.1002/jps.10094.

(32) Wilding, K. M.; Zhao, E. L.; Earl, C. C.; Bundy, B. C. Thermostable Lyoprotectant-Enhanced Cell-Free Protein Synthesis for on-Demand Endotoxin-Free Therapeutic Production. New Biotechnology 2019, 53, 73–80. 10.1016/j.nbt.2019.07.004.

(33) Izutsu, K.; Yoshioka, S.; Terao, T. Stabilizing Effect of Amphiphilic Excipients on the Freeze-thawing and Freeze-drying of Lactate Dehydrogenase. Biotech & Bioengineering 1994, 43 (11), 1102–1107. 10.1002/bit.260431114.

(34) Warfel, K. F.; Williams, A.; Wong, D. A.; Sobol, S. E.; Desai, P.; Li, J.; Chang, Y.-F.; DeLisa, M. P.; Karim, A. S.; Jewett, M. C. A Low-Cost, Thermostable, Cell-Free Protein Synthesis Platform for On-Demand Production of Conjugate Vaccines. ACS Synth. Biol. 2023, 12 (1), 95–107. 10.1021/acssynbio.2c00392.

(35) Lu, Y. Textile-Embedded Cell-Free Biosensors. Nat. Biomed. Eng 2022, 6 (3), 225–226. 10.1038/s41551-022-00869-3.

(36) Nguyen, P. Q.; Soenksen, L. R.; Donghia, N. M.; Angenent-Mari, N. M.; De Puig, H.; Huang, A.; Lee, R.; Slomovic, S.; Galbersanini, T.; Lansberry, G.; Sallum, H. M.; Zhao, E. M.; Niemi, J. B.; Collins, J. J. Wearable Materials with Embedded Synthetic Biology Sensors for Biomolecule Detection. Nat Biotechnol 2021, 39 (11), 1366–1374. 10.1038/s41587-021-00950-3.

